# Hearing loss promotes schizophrenia-relevant brain and behavioral abnormalities in a mouse model of human 22q11.2 Deletion Syndrome

**DOI:** 10.1101/539650

**Authors:** Fhatarah A. Zinnamon, Freya G. Harrison, Sandra S. Wenas, Arne F. Meyer, Qing Liu, Kuan Hong Wang, Jennifer F. Linden

## Abstract

Hearing loss has been implicated as a risk factor for schizophrenia, but it is not known whether this association arises from common etiology, top-down influences (e.g., social isolation), bottom-up neurobiological mechanisms, or combinations of these factors. Patients with 22q11.2 Deletion Syndrome (22q11.2DS) have a 25-30% risk of developing schizophrenia, and also suffer frequent hearing loss. Here, we used the *Df1*/+ mouse model of 22q11.2DS to investigate the relationship between hearing loss and susceptibility to schizophrenia-relevant brain and behavioral abnormalities. *Df1*/+ mice have a multi-gene deletion analogous to the chromosomal microdeletion that causes human 22q11.2DS, and like human 22q11.2DS patients exhibit high rates of hearing loss arising primarily from susceptibility to middle ear inflammation. We found that hearing loss in *Df1*/+ mice affected schizophrenia-relevant endophenotypes, including electrophysiological measures of central auditory gain and behavioral measures of auditory sensorimotor gating. Moreover, PV+ inhibitory interneurons, another marker for schizophrenia pathology, were significantly reduced in density in auditory cortex but not secondary motor cortex of *Df1*/+ mice with hearing loss. These results reveal bottom-up neurobiological mechanisms through which peripheral hearing loss arising from the 22q11.2 deletion may promote the emergence of schizophrenia-relevant auditory brain and behavioral abnormalities, and also suggest a link between conductive hearing loss and reduced PV+ interneuron density in the auditory cortex.

**SIGNIFICANCE STATEMENT:** Hearing loss is a known risk factor for schizophrenia. Deletion of chromosomal locus 22q11.2 is associated with both schizophrenia and hearing loss in humans. In the *Df1*/+ mouse model of human 22q11.2 Deletion Syndrome, we find that hearing loss shapes measures that are considered schizophrenia-relevant endophenotypes, such as central auditory gain and auditory sensorimotor gating. Moreover, we report a reduction in density of PV+ inhibitory interneurons in the auditory cortex, but not secondary motor cortex, of *Df1*/+ mice with hearing loss. These results suggest mechanisms through which hearing loss associated with the 22q11.2 deletion may promote emergence of schizophrenia-relevant auditory brain and behavioral abnormalities and indicate that conductive hearing loss may influence PV+ interneuron density in the auditory cortex.

## INTRODUCTION

22q11.2 Deletion Syndrome (22q11.2DS) is the strongest known cytogenetic risk factor for schizophrenia in humans (McDonald-McGinn et al., 2015). 22q11.2DS is caused by a 0.7-3Mbp congenital multigene deletion which occurs in approximately 1 out of 3000-6000 live births, making it the most common chromosomal microdeletion syndrome and the second most common chromosomal disorder after Down’s Syndrome (McDonald-McGinn et al., 2015; Paylor and Lindsay, 2006). Approximately 25-30% of patients with 22q11.2DS develop schizophrenia during adolescence or adulthood (Drew et al., 2011; McDonald-McGinn et al., 2015). In addition to increased schizophrenia susceptibility, 22q11.2DS is also associated with over 100 different malformations and clinical presentations including heart defects, immune dysfunction, hypocalcaemia, and craniofacial abnormalities such as cleft palate (Paylor and Lindsay, 2006). Furthermore, patients with 22q11.2DS have frequent hearing loss, arising primarily from high rates of recurrent or chronic otitis media (middle ear inflammation) (Verheij et al., 2017).

The *Df1*/+ mouse model of 22q11.2DS has an engineered hemizygous deletion of 1.2Mbp encompassing 18 orthologs of genes deleted in human 22q11.2DS (Lindsay et al., 1999). Along with other mouse models of 22q11.2DS, the *Df1*/+ mouse recapitulates many phenotypic features of human 22q11.2DS (Paylor and Lindsay, 2006), including cardiac defects (Lindsay et al., 1999), craniofacial abnormalities (Aggarwal et al., 2006), and brain and behavioral abnormalities that have been linked to schizophrenia in humans (e.g., Hamm, Peterka, Gogos, & Yuste, 2017; Sigurdsson, Stark, Karayiorgou, Gogos, & Gordon, 2010). Behaviorally, *Df1*/+ mice exhibit reduced prepulse inhibition of the acoustic startle response (Paylor et al., 2001), a robust behavioral marker for schizophrenia-like abnormalities and common feature of 22q11.2DS in humans (Drew et al., 2011). Functionally, specific abnormalities in auditory thalamocortical processing have also been reported in *Df1*/+ mice, including abnormal sensitivity of auditory thalamocortical projections to antipsychotic drugs (Chun et al., 2014). Interestingly, like humans with 22q11.2DS, *Df1*/+ mice have been found to have high rates of hearing loss (Fuchs et al., 2013), due to increased susceptibility to otitis media caused by developmental defects arising from the 22q11.2DS deletion (Fuchs et al., 2015). However, the potential interaction between hearing loss and auditory brain and behavioral abnormalities in *Df1*/+ mice has not previously been explored.

In humans, hearing loss has been described as the “neglected risk factor for psychosis” (Sommer et al., 2014). Hearing loss is associated with increased risk of psychosis and hallucinations, and hearing impairment in childhood elevates the risk of developing schizophrenia later in life (see Linszen, Brouwer, Heringa, & Sommer, 2016). The mechanisms underlying the association between hearing loss and schizophrenia are unknown, and could include common etiology, top-down influences (e.g., social isolation), and/or bottom-up effects. A role for bottom-up effects is suggested by data from animal studies indicating that loss of peripheral auditory input drives long-lasting changes in central auditory processing which can affect behavior (Chambers et al., 2016; Sanes and Kotak, 2011; Takesian et al., 2009; Yao and Sanes, 2018). Even reversible conductive hearing loss, such as that caused by otitis media, can produce persistent changes in inhibitory synaptic transmission in the auditory cortex (Mowery et al., 2015; Sanes and Kotak, 2011; Takesian et al., 2012, 2009). However, it is not yet known how changes in the auditory brain arising from hearing loss relate to changes in brain circuitry produced by genetic and other risk factors for schizophrenia. Parvalbumin-positive (PV+) interneurons play a key role in maintaining excitation-inhibition balance in the cortex (Moore et al., 2018) and abnormalities in these inhibitory cells are thought to be important to the pathophysiology of schizophrenia (Lewis, 2014). However, reductions in the density of PV+ interneurons have not previously been linked with conductive hearing loss in animal models of either schizophrenia or otitis media.

Here we report a link between hearing loss and schizophrenia-relevant auditory brain and behavioral abnormalities in the *Df1*/+ mouse model of 22q11.2DS. Putative schizophrenia endophenotypes in *Df1*/+ mice, such as abnormalities in electrophysiological markers of central auditory gain and behavioral measures of auditory sensorimotor gating, were largely explained by hearing loss. Moreover, density of PV+ interneurons was significantly reduced in the auditory cortex of *Df1*/+ mice with hearing loss. Our results suggest that peripheral hearing loss arising from the 22q11.2 deletion contributes to the emergence of schizophrenia-relevant auditory brain and behavioral abnormalities in 22q11.2DS.

## MATERIALS AND METHODS

### Animals

Experiments were conducted in *Df1*/+ mice and their wild-type (WT) littermates, all 6 to 19 weeks of age (mean 13, SD 4.49 weeks). All mice used in these experiments were bred from a genetically modified line established previously on the C57BL/6J background (Lindsay et al., 1999). Breeding was maintained by crossing *Df1*/+ males from within the colony with WT C57BL/6J females either from within the colony or from Charles River UK. Back-crossing into the C57BL/6J strain had been maintained for well over 10 generations at the start of the experiments. Mice were maintained in standard cages and mouse housing facilities, on a standard 12 h on, 12 h off light/dark cycle. All experiments were performed in accordance with a Home Office project license approved under the United Kingdom Animal Scientific Procedures Act of 1986.

### Acoustic startle response (ASR) and prepulse inhibition (PPI) testing

To test auditory sensorimotor gating in *Df1*/+ and WT mice, we measured the acoustic startle response (ASR) and prepulse inhibition (PPI) of the ASR. Startle experiments were conducted using a custom-built startle measurement system consisting of four piezoelectric sensors mounted underneath a circular platform 12.5 cm in diameter, housed within a dark acoustic isolation booth lined with acoustical foam. The platform was encircled by a removable cylindrical plastic tube approximately 13 cm in diameter and 30 cm high, to prevent the mice from jumping off. Infrared lighting outside the visible range of the mouse and a webcam (Logicam) were used to monitor the animals during testing. If a mouse tried to climb up the cylinder, the testing session was paused, the animal was returned to the platform and the recording restarted. A speaker (Peerless Vifa XT25TG30-04) was mounted in the ceiling of the booth, approximately 40 cm above the platform and centered over the opening of the cylindrical platform enclosure.

Sensor signals were acquired at a 30 kHz sample rate from the analog inputs on an Open Ephys data acquisition board using Open Ephys GUI software (www.open-ephys.org), then downsampled to 1 kHz for analysis. Acoustic stimuli were generated at a 96000 Hz sample rate using a professional sound card (RME HDSPe AIO) and custom Python software (“StimControl”, by Arne F. Meyer). Stimuli were delivered to the speaker via either a Rotel RB-971 speaker amplifier or a Tucker-Davis Technologies SA5 speaker amplifier. Speaker output was calibrated using a ¼-inch free-field microphone, preamplifer and amplifier (G.R.A.S. 40BF, 26AC and 12AA) before the experiments began, to ensure that noise levels for cue and startle stimuli were within 2 dB of target levels. Measurements of speaker output were made with the microphone positioned in the center of the platform at the approximate height of the animals’ ears during testing.

Acoustic startle and prepulse inhibition data were typically collected from each mouse in two repeated 25-30 min testing sessions on different days, separated by 2 days. An additional testing session was conducted on a third day if a previous session had been unsuccessful (e.g., if there had been technical problems or difficulty keeping the animal on the measurement platform). Each session involved two types of behavioral tests: startleIO tests and startlePPI tests.

In startleIO tests, the ASR input/output function was measured, recorded, and also visually inspected online to choose prepulse cue levels used for subsequent startlePPI tests. StartleIO tests involved presentation of a 65 dB SPL continuous white noise background sound, overlaid with 20 ms noise bursts with sound levels ranging from 55 or 60 dB SPL to 95 dB SPL in 5 dB steps. Noise bursts of different sound levels were presented in pseudorandom order, with randomly varied inter-burst intervals that typically ranged from 1.67 s to 8 s (average 5 s). A typical startleIO test lasted 225 s and involved presentation of 5 repeats of each noise burst sound level. An initial online analysis of startle magnitude as a function of noise burst intensity level was performed to choose the two or three loudest sound levels that did not evoke a startle response, for use in subsequent startlePPI tests.

In startlePPI tests, prepulse inhibition of the acoustic startle response was assessed. Like startleIO tests, startlePPI tests were conducted with a continuous 65 dB SPL white noise background. Approximately 20-25% of startlePPI trials were uncued trials, in which a 40 ms, 95 dB SPL startle-eliciting stimulus was presented in the middle of the 20 s trial. On the remaining 75-80% of trials (cued trials), the startle-eliciting stimulus was preceded by a 20 ms prepulse noise burst at one of the sound levels chosen to be below the startle threshold in the startleIO tests. The delay between the onset of the prepulse cue and the onset of the startle-eliciting stimulus alternated randomly between 50 ms and 100 ms; however, PPI proved to be more robust in both *Df1*/+ and WT mice using the shorter delay, and so only data collected using the 50 ms delay is presented here. A typical startlePPI test lasted 1000 s and involved pseudorandom presentations of 10 repeats of each of 4 possible cue conditions (2 sound levels × 2 delays) and 10 repeats of the uncued condition.

Within 1-2 days of the final behavioral testing session, mice that underwent startle testing were anaesthetized for auditory brainstem response and auditory evoked potential recordings, and then immediately terminated and perfused for immunohistochemical analysis.

### Auditory brainstem response (ABR) and auditory evoked potential (AEP) recording

To screen mice for hearing loss we recorded the auditory brainstem response (ABR), an electroencephalographic signal measured through the use of scalp electrodes which allows for the detection and visualization of sound-evoked potentials generated by neuronal circuits in the ascending auditory pathway (Willott, Carlson, and Chen 1994). ABR recordings were performed in a sound isolation booth (Industrial Acoustics Company, Inc.). Auditory stimuli were generated at a sample rate of 195,312.5 Hz using a digital signal processor (Tucker-Davis Technologies, TDT RX6), attenuated as needed (TDT PA5), amplified (TDT SA1), and presented using a free-field speaker (TDT FF1) positioned 17-18 cm from the ear directed toward the speaker. Speaker output was calibrated to within 2 dB of target values before each set of experiments using a Bruel & Kjaer ¼ inch microphone (4939), placed at the location of the ear to be tested. Data was acquired at a 24,414 Hz sample rate (TDT RX5) using a low impedance headstage and signal amplifier (TDT RA4LI and RA16SD, 20x gain overall, 2.2 Hz - 7.5kHz filtering) along with a custom low-pass filter designed to remove attenuation switching transients (100 kHz cutoff). Stimulus presentation and data acquisition was controlled using software from TDT (Brainware) and custom software written in MATLAB (Mathworks).

Mice were anaesthetized via intraperitoneal injection of a ketamine-medetomidine cocktail (0.003-0.01 ml/g body weight of 10mg/ml ketamine, 0.083 mg/ml metetomidine). Body temperature was maintained at 37-38°C using a homeothermic blanket (Harvard Apparatus). Subdermal needle electrodes (Rochester Medical) were typically inserted under the skin at the vertex (positive), at the bulla behind the ear directed toward the speaker (negative), and over the olfactory bulb (ground). In a few animals for which ABRs but not AEPs were recorded, we instead placed the ground electrode behind the bulla opposite the tested ear. ABR thresholds were determined in the left and right ears in turn in most animals. The animal was re-oriented between recordings to direct the ear being tested toward the speaker, but we avoided altering the positions of the subdermal electrodes by switching electrodes at the input of the pre-amplifier whenever possible. ABR stimuli were 50 μs monophasic clicks ranging in sound level from 0 to 90 dB SPL in 5 dB steps, repeated 500 times at an inter-click interval of 50 ms.

In a subset of animals, we collected further ABR recordings in combination with auditory cortical evoked-potential (AEP) recordings. To measure AEP signals, additional subdermal electrodes were placed at locations corresponding approximately to the left and right auditory cortices, and referenced to the same ground (at the olfactory bulb) as for the ABR recordings. Stimuli were 80 dB SPL clicks presented 1000 times at an inter-click interval of 300 ms. The longer inter-click interval allowed for resolution of late cortical AEP waves as well as the earlier ABR signals.

### Histology, immunohistochemistry and microscopy

Mice were euthanized using an overdose of sodium pentobarbital (0.1-0.2 ml of 20 mg/ml Euthatal; Rhône Mérieux, Essex, UK) and were perfused transcardially with at least 30 ml 4% paraformaldehyde (Merck, Dorset, UK) using a peristaltic pump. Brain tissue was removed and stored in 4% paraformaldehyde at 4°C until histology could be performed. Brains were transferred to increasing concentrations of sucrose solution (15% followed by 30%) for cryoprotection before freezing and slicing. Coronal sections 50 μm thick were cut using a cryostat (Bright Instruments). In some animals, alternate sections were used for Nissl staining and parvalbumin (PV) immunohistochemistry. In other animals, sections were triple-stained for PV, NeuN and DAPI. Nissl staining was performed on mounted sections using cresyl violet, according to standard procedures. For PV and NeuN staining, sections were incubated with 0.5% Triton and blocked with 0.5% Triton and goat serum blocking solution before being incubated in mouse anti-PV monoclonal antibody (1:2000, Sigma-Aldrich) and mouse anti-NeuN polyclonal antibody (1:2000, Sigma-Aldrich) at 4°C overnight. Sections were then rinsed in PBS and incubated with goat anti-mouse secondary antibody (1:200, Sigma-Aldrich) for one hour at room temperature on a shaker. Sections were then rinsed again in PBS. Slides were mounted with DAPI Flouromount-G (Southernbiotech) and coverslipped.

Sections stained for Nissl were viewed at 2.5× to 5× magnifications on a Zeiss Axio Scan 2 Imaging microscope. Single-plane images of appropriate A1 and M2 sections were taken at 5× to 10× magnification, with individual images for each hemisphere. Sections immunostained for PV were viewed and captured at 10× magnification on a Zeiss Axio Scan 2 Imaging microscope. All images were obtained at a resolution of 720 pixels/inch.

### Data analysis

A total of 32 *Df1*/+ mice (11 male, 21 female) and 39 WT littermates (20 male, 19 female) were used in these experiments. All data analyses were conducted blind to the genotype of the animal. We performed comparisons between two groups with the unpaired t-test, and comparisons between multiple groups with the ANOVA followed by Fisher’s LSD tests where group differences were significant. Non-parametric tests were used where required; for random permutation tests the data were randomly permuted 10,000 times and the actual difference between the means of the groups were compared against the randomized distribution. Unless otherwise indicated, all statistical tests were two-tailed with α=0.05.

### Startle data analysis

Startle analysis was performed on data from 17 *Df1*/+ mice and 14 WT mice that completed startle testing successfully and also underwent subsequent ABR testing to assess hearing sensitivity. The time-varying startle signal was defined at each 1-ms time point as the square root of the sum of squared outputs from the four piezoelectric force sensors embedded beneath the startle platform. The startle response was defined as the sum of these sample-by-sample startle signals over the 0-65 ms period following the onset of the startle-eliciting noise burst. To determine the threshold for the startle response for each mouse, data from startleIO behavioral sessions were pooled across repeated sessions. The startle response threshold was then defined as the lowest noise burst level for which its startle response was significantly greater than the startle response elicited by a 60 dB SPL noise burst (Wilcoxon rank-sum test, p<0.01), and for which all noise burst levels above the threshold sound level also elicited a significantly stronger startle response than the 60 dB SPL reference. Mice for which no startle response threshold could be identified (2 *Df1*/+, 5 WT) were excluded from the population analysis shown in Figure 8.

The uncued startle response was determined using data from startlePPI behavioral sessions, and computed as the mean startle response to a 95 dB SPL noise burst on uncued trials from each behavioral session, averaged across repeated behavioral sessions for each animal. A behavioral session was included in this and other PPI analyses only if the uncued startle response was significantly different from the background startle response computed over a period 0–65 ms before the noise burst occurred (Wilcoxon sign-rank test, p<0.01). For startlePPI sessions meeting this criterion, the cued startle response was then computed for each cue type, as the mean startle response to the 95 dB SPL noise burst on cued trials in each behavioral session averaged across repeated behavioral sessions for each animal. Prepulse inhibition (PPI) was then defined for each behavioral session and each cue type as follows:

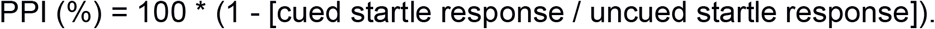

For population analysis, PPI values for each cue type and each behavioral session were averaged across behavioral sessions to obtain a single PPI measure for each animal. Cue conditions in which the cue itself evoked a significant startle response (i.e., significant difference in background startle response 0–65 ms before startle noise onset between cued and uncued trials; Wilcoxon rank-sum test, p<0.01) were excluded from the population analysis.

### ABR/AEP data analysis

ABR recordings were performed in 25 *Df1*/+ (9 male, 16 female) and 26 WT (13 male, 13 female) mice. A total of 104 ABR threshold measurements (50 *Df1*/+, 52 WT) were obtained; each mouse contributed two separate ABR measurements, one for each ear. In the majority of animals, we collected ABR recordings in combination with auditory cortical evoked-potential (AEP) recordings. Combined ABR/AEP recordings were obtained from 79 ears and contralateral hemispheres (40 *Df1*/+, 39 WT); 20 *Df1*/+ mice and 19 WT mice each contributed separate ABR/AEP measurements for both of the two possible ear / contralateral hemisphere combinations, while 1 WT mice died after only one ABR/AEP recording had been completed. Separate measurements from the same animal obtained during auditory stimulation of left versus right ears were treated as independent measurements in our data analysis.

The differential ABR signal for a given ear (vertex electrode signal minus ipsilateral ear electrode signal) was averaged across 500 trials for each click intensity, and analyzed to extract ABR waveform components as in previous studies (Fuchs et al. 2013). The ABR threshold was defined as the lowest sound intensity at which at least one of the characteristic deflections in the ABR waveform could be distinguished upon visualization of the ABR signal across all sound intensity levels (see Fuchs et al., 2013 for examples and details).

The single-ended AEP signal was derived from the temporal lobe electrode contralateral to the ear directed at the speaker, since AEP signals were stronger over the contralateral than ipsilateral hemisphere. Signals were averaged over 1000 repeated trials to obtain the AEP waveform. The AEP wave components were characterized as in Maxwell et al., 2004: P1, or P20, was defined as the maximum peak 15-30 ms after stimulus onset; N1, or N40, was defined as the maximum negative deflection 25-60 ms post-onset; and P2, or P80, was defined as the maximum peak 60-110 ms post-onset.

Wave amplitudes were defined for an 80 dB SPL click stimulus, as the difference in ABR/AEP signal amplitudes between baseline at the start of the recording and peak for ABR wave I; between the peak and subsequent trough for AEP P1-N1 complex; and between the trough and subsequent peak for AEP N1-P2 complex (see Figure 2). Wave latencies were defined as peak or trough times relative to stimulus onset. All amplitude and latency analyses were performed blind to mouse genotype and ABR threshold. Central auditory gain was defined as the amplitude of the AEP wave complex (P1-N1 or N1-P2) divided by the amplitude of ABR wave I.

### Quantification of cell counts

PV+ and NeuN+ cell counts and laminar distributions in A1 and M2 were estimated for images from both hemispheres in each mouse when possible. Immunohistochemical data from some hemispheres and animals were lost due to problems with perfusion, damage to sections, or aberrant fluorescence in images. See Table 1 for details of number of mice and hemispheres used for different comparisons shown in Figure 8, Figure 9 and Figure 10. Only data from animals that had undergone ABR testing prior to immunohistochemistry were used for comparisons between *Df1*/+ mice with hearing loss, *Df1*/+ mice without hearing loss, and WT mice.

**Table 1.**
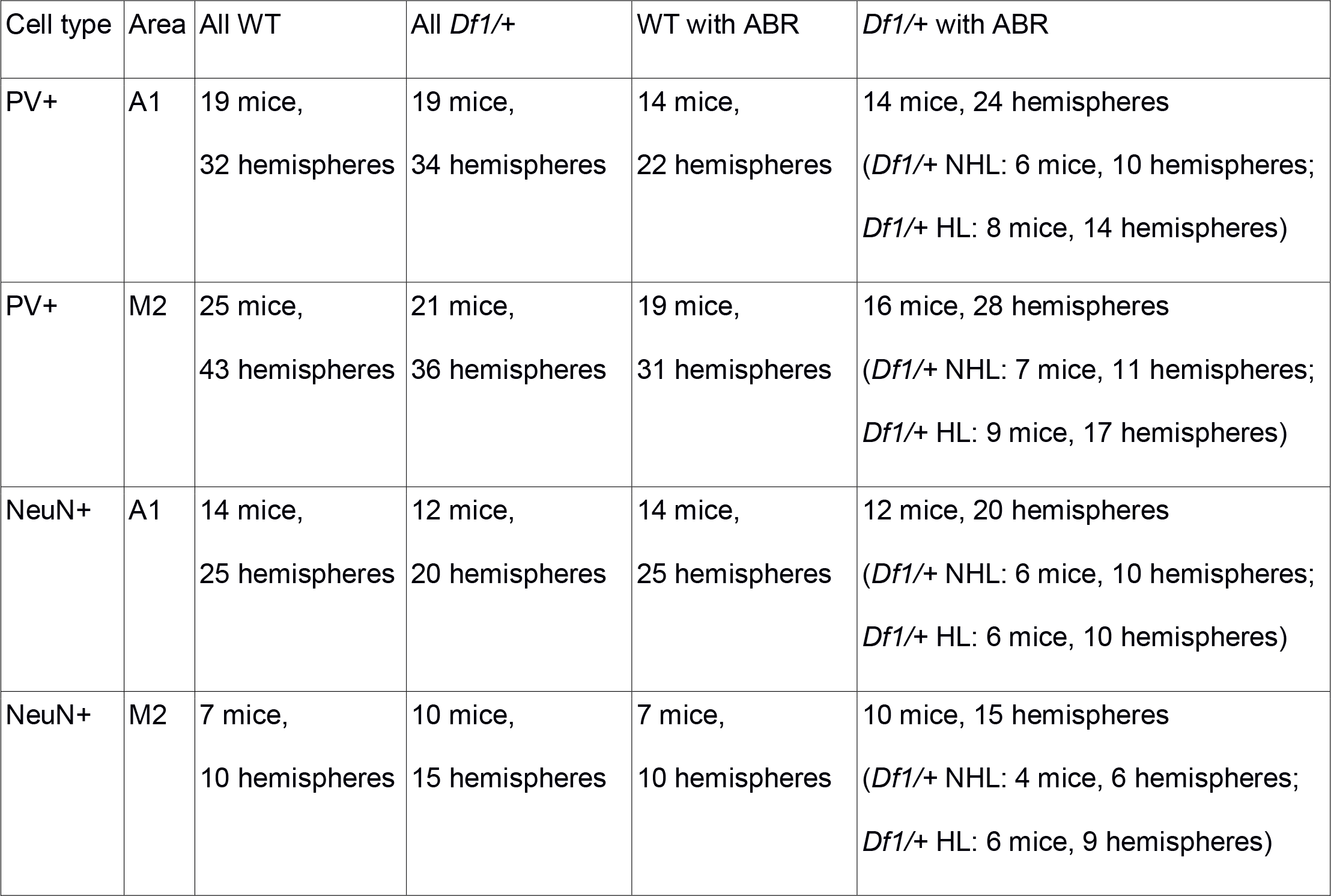
Numbers of mice and brain hemispheres used for analyses of PV+ and NeuN+ cell density and laminar distribution. All mice were included in comparisons of WT and *Df1*/+ mice (Figure 8B,E, Figure 9B,E and Figure 10A,C). Only data from mice which underwent ABR testing were included in comparisons of WT animals and *Df1*/+ mice with and without hearing loss (Figure 8C,F, Figure 9C,F and Figure 10B,D).

| Cell type | Area | All WT | All *Df1/+* | WT with ABR | *Df1/+* with ABR |
| --- | --- | --- | --- | --- | --- |
| PV+ | A1 | 19 mice,<br>32 hemispheres | 19 mice,<br>34 hemispheres | 14 mice,<br>22 hemispheres | 14 mice, 24 hemispheres<br>(*Df1/+* NHL: 6 mice, 10 hemispheres;<br>*Df1/+* HL: 8 mice, 14 hemispheres) |
| PV+ | M2 | 25 mice,<br>43 hemispheres | 21 mice,<br>36 hemispheres | 19 mice,<br>31 hemispheres | 16 mice, 28 hemispheres<br>(*Df1/+* NHL: 7 mice, 11 hemispheres;<br>*Df1/+* HL: 9 mice, 17 hemispheres) |
| NeuN+ | A1 | 14 mice,<br>25 hemispheres | 12 mice,<br>20 hemispheres | 14 mice,<br>25 hemispheres | 12 mice, 20 hemispheres<br>(*Df1/+* NHL: 6 mice, 10 hemispheres;<br>*Df1/+* HL: 6 mice, 10 hemispheres) |
| NeuN+ | M2 | 7 mice,<br>10 hemispheres | 10 mice,<br>15 hemispheres | 7 mice,<br>10 hemispheres | 10 mice, 15 hemispheres<br>(*Df1/+* NHL: 4 mice, 6 hemispheres;<br>*Df1/+* HL: 6 mice, 9 hemispheres) |

Cortical areas of interest were defined by overlaying the section image with the Paxinos and Franklin (2012) mouse atlas image of the corresponding coronal location relative to bregma, using Adobe Photoshop Elements (Adobe Systems Inc). Using ImageJ, dots were centered over the cells, cell centroid locations recorded (X,Y coordinates, in pixels), and data transferred to MATLAB for analysis of cell counts. All cells in the cortical area of interest (A1 or M2) were counted in sections immunolabelled for PV, and cells within a 5% area through the center of the region of interest were counted in sections immunolabelled for NeuN. Laminar distributions of cells were estimated using custom MATLAB software designed to calculate cell centroid depth along a line perpendicular to the pial surface and white matter. Laminar distributions were normalized by the pia-to-white-matter distance in each section. Laminar distributions were compared between animal groups using multiple t-tests on cell counts binned by cortical depth; similar results were obtained using 5, 10 or 20 bins in depth. All cell counts and laminar distribution data were gathered blind to genotype and ABR thresholds of the mouse.

## RESULTS

### *Df1*/+ mice exhibit hearing loss in 60% of animals (54% of ears)

Both human 22q11.2DS patients and mouse models of 22q11.2DS have been reported to have high rates of hearing loss due primarily to middle ear inflammation (Dyce et al., 2002; Fuchs et al., 2015, 2013; Reyes et al., 1999; Verheij et al., 2017). We measured hearing thresholds for each ear in 25 *Df1*/+ mice and 26 WT mice using the auditory brainstem response (ABR). The ABR is an electroencephalographic signal arising from sound-evoked neural activity within the auditory nerve and brainstem (Willott, 2006). The ABR threshold was defined as the lowest sound intensity at which at least one of the characteristic deflections in the ABR waveform could be distinguished upon visualization of the signal across all sound intensity levels (see Materials and Methods). Click-evoked ABR thresholds were estimated for each ear in each mouse by an observer blind to the genotype of the animal. To minimize the influence of age-related sensorineural hearing loss characteristic of the C57BL/6J background strain (Parham, 1997), *Df1*/+ and WT mice tested in each session were littermates between 6 and 12 weeks of age. To reduce the potential for transmission of sound signals to the inner ear via bone or tissue conduction rather than via the middle ear, all sound stimuli were presented free-field rather than through an in-ear coupler.

Replicating previous results obtained in a different cohort of *Df1*/+ and WT mice (Fuchs et al. 2013), we found clear evidence for hearing loss in more than half of the *Df1*/+ animals (Figure 1). Mean ± SEM values for ABR thresholds were 44.8 ± 1.73 dB SPL in *Df1*/+ ears versus 32.5 ± 0.47 dB SPL in WT ears. Elevation of ABR thresholds in *Df1*/+ relative to WT mice was evident in both male and female animals (Figure 1A), and a two-way ANOVA revealed a significant effect of genotype but not gender (genotype: F(1,97)=44.24, p<0.0001; gender: F(1,97)=0.63, p=0.43). The range of ABR thresholds in *Df1*/+ ears extended from the minimum to well beyond the maximum of the WT range (Figure 1B); i.e., ABR thresholds were abnormally elevated in some *Df1*/+ ears but not others. Defining the upper bound of normal hearing as 2.5 SD above the mean ABR threshold for WT ears (40.89 dB SPL), we found that 54% (27/50) of *Df1*/+ ears and 0% (0/52) of WT ears displayed hearing loss. Overall, 60% (15/25) of *Df1*/+ mice had either monaural or binaural hearing loss, and monaural hearing loss occurred most commonly in the left ear (Figure 1C). These results align both qualitatively and quantitatively with findings previously reported by Fuchs et al. (2013).

**Figure 1.**
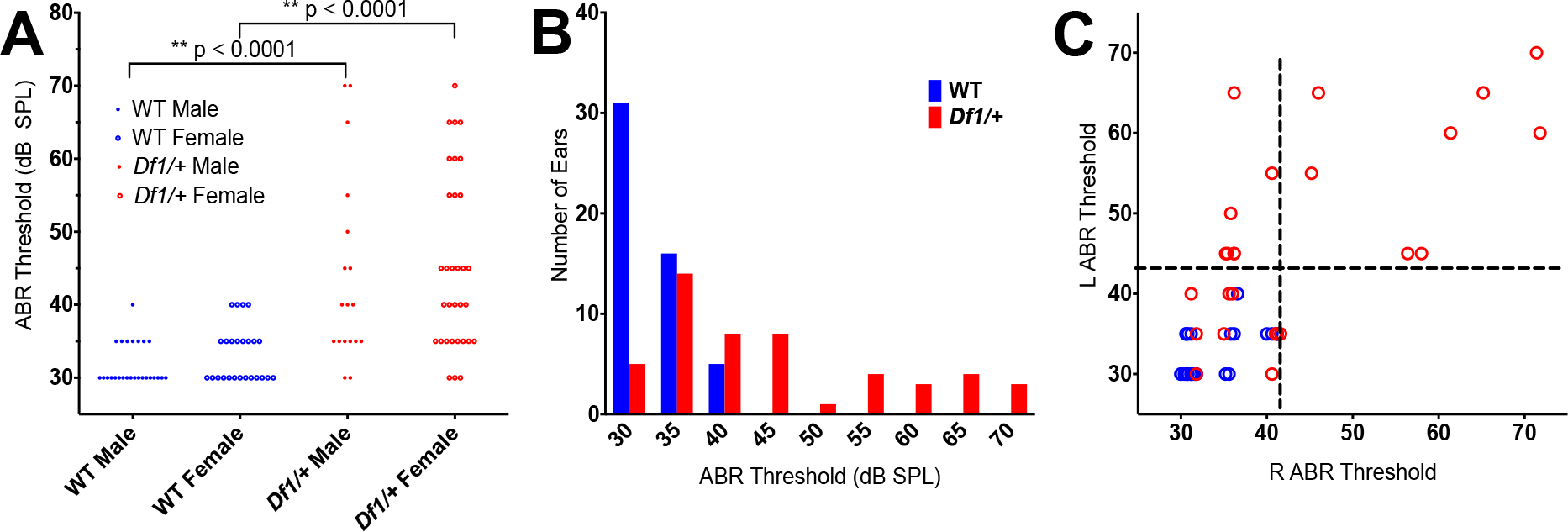
Elevated auditory brainstem response (ABR) thresholds in *Df1*/+ mice. (A) Click ABR thresholds recorded from individual ears in male and female WT (blue) and *Df1*/+ (red) mice. Two-way ANOVA with genotype and gender as independent variables reveals a significant effect of genotype (F(1,97)=44.24, p<0.0001) but not of gender (F(1,97)=0.63, p=0.43). Data points represent individual ear measurements; each animal contributed two measurements, one for each ear. Number of mice: 13 WT male, 13 WT female, 9 *Df1*/+ male, 16 *Df1*/+ female. (B) Click ABR thresholds pooled across recordings from male and female animals. Note that the *Df1*/+ ABR threshold distribution extends from the minimum to well beyond the maximum of the WT range. (C) Scatter plot showing the relationship between left and right ear ABR thresholds in each mouse. Dashed lines indicate the upper bound of normal hearing, defined as 2.5 SD above the mean ABR threshold for WT ears (see text). 60% (15/25) of *Df1*/+ mice had either monaural or binaural hearing loss, and 54% (27/50) of *Df1*/+ ears exhibited hearing loss. As previously observed (Fuchs et al., 2013), monaural hearing loss in *Df1*/+ mice occurred most commonly in the left ear (cf. Figure 2 in Fuchs et al., 2013).

### ABR Wave I amplitude reductions in *Df1*/+ mice with hearing loss are not maintained in cortical AEPs

Previous studies have reported abnormalities in sound-evoked auditory thalamic and/or cortical activity in mouse models of 22q11.2DS (Chun et al., 2014; Didriksen et al., 2017). We wondered if auditory brain responses might differ between *Df1*/+ and WT mice, and if so, whether these differences might be related to peripheral hearing loss. To find out, we recorded both ABRs and auditory evoked potentials (AEPs) in 20 WT and 20 *Df1*/+ mice (11 *Df1*/+ with no hearing loss, 9 *Df1*/+ with hearing loss in at least one ear), following presentations of loud (80 dB SPL) clicks at 300 ms inter-click intervals. We then measured latencies and amplitudes of ABR and AEP waves to determine whether the waveforms differed between WT and *Df1*/+ mice, or between WT mice and *Df1*/+ mice with and without hearing loss (Figure 2, Figure 3, and Figure 4). To assess the timing and strength of afferent input to the auditory brain, we measured the peak latency and baseline-to-peak amplitude of ABR wave I (Figure 2A), which arises from the auditory nerve (Willott, 2006). Within the AEP, we focused on wave peaks or troughs typically attributed to activity within the auditory thalamus (P1), auditory cortex (N1), and associative cortices (P2), measuring P1, N1, and P2 latency and P1-N1 and N1-P2 amplitudes (Figure 2D).

**Figure 2.**
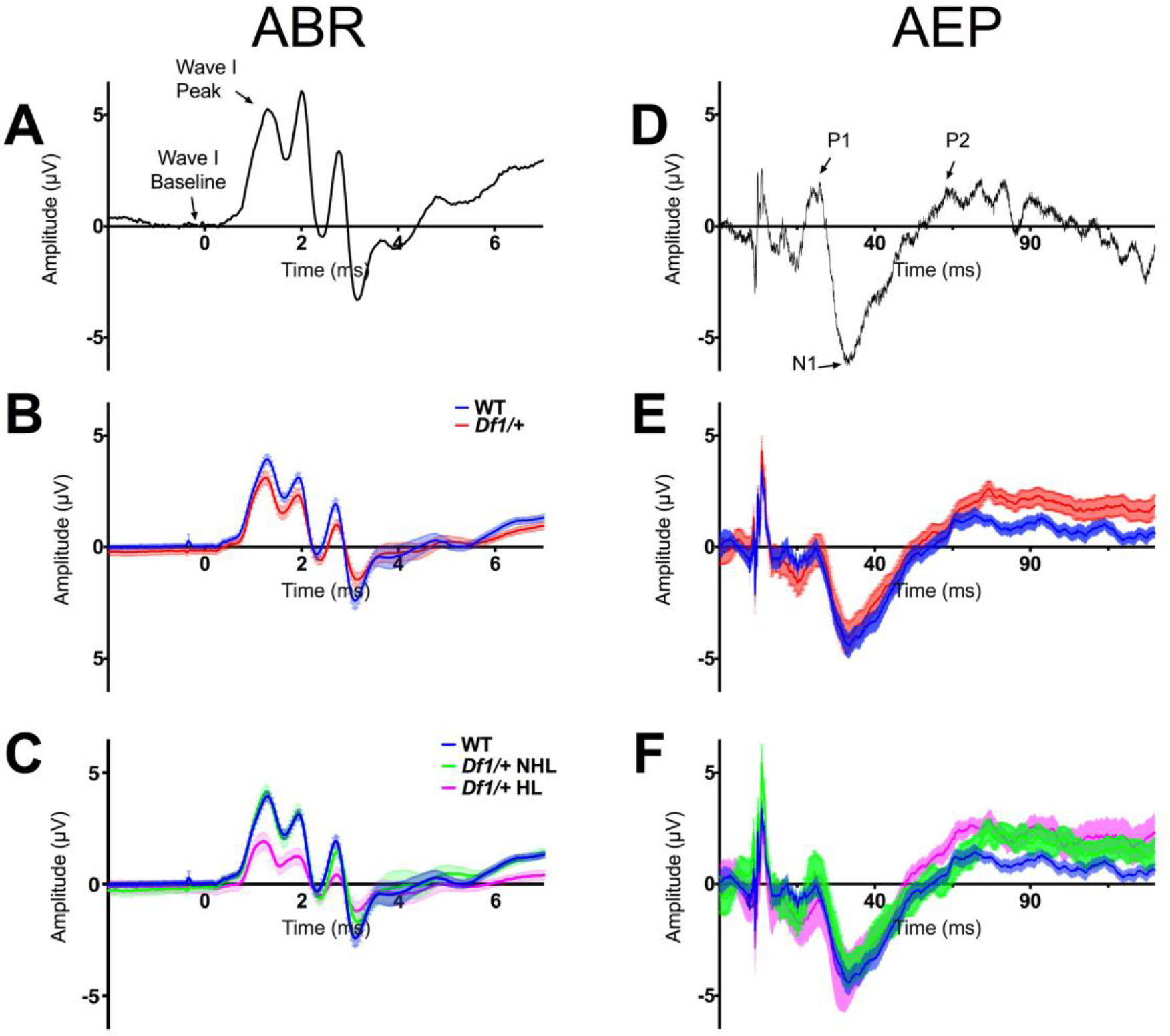
Average ABR and auditory evoked potential (AEP) waveforms in WT and *Df1*/+mice. (A) Example trial-averaged ABR to an 80 dB SPL click, recorded ipsilateral to the stimulated ear in an individual WT animal. Arrows indicate baseline and peak used for measurement of wave I amplitude. (B) Mean ABR waveforms averaged across recordings from WT mice (blue) and *Df1*/+ mice (red). Error bars indicate SEM across all trial-averaged ABR recordings for each group of animals. (C) Same as B, but with *Df1*/+ ABR recordings separated into those from *Df1*/+ mice without hearing loss (green) or *Df1*/+ mice with hearing loss in at least one ear (magenta). (D) Example trial-averaged AEP to an 80 dB SPL click, recorded over auditory cortex contralateral to the stimulated ear in a WT animal. Arrows indicate P1 peak, N1 trough, and P2 peak. (E) Mean AEP waveforms averaged across recordings from different mice; color conventions as in B. Error bars indicate SEM across all trial-averaged AEP recordings for each group of animals. (F) Same as E, but with *Df1*/+ AEP recordings separated into those from *Df1*/+ mice with or without hearing loss in at least one ear; color conventions as in C. Ipsilateral ABR and contralateral AEP data were obtained from the same recording for each stimulated ear; however ABR waveforms in A-C represent differential signals while AEP waveforms in D-F are single-ended signals (see Materials and Methods). Number of mice: 20 WT, 20 *Df1*/+ (11 *Df1*/+ without hearing loss, 9 *Df1*/+ with hearing loss). Number of ABR/AEP recordings: 39 WT, 40 *Df1*/+ (22 *Df1*/+ without hearing loss, 18 *Df1*/+ with hearing loss).

**Figure 3.**
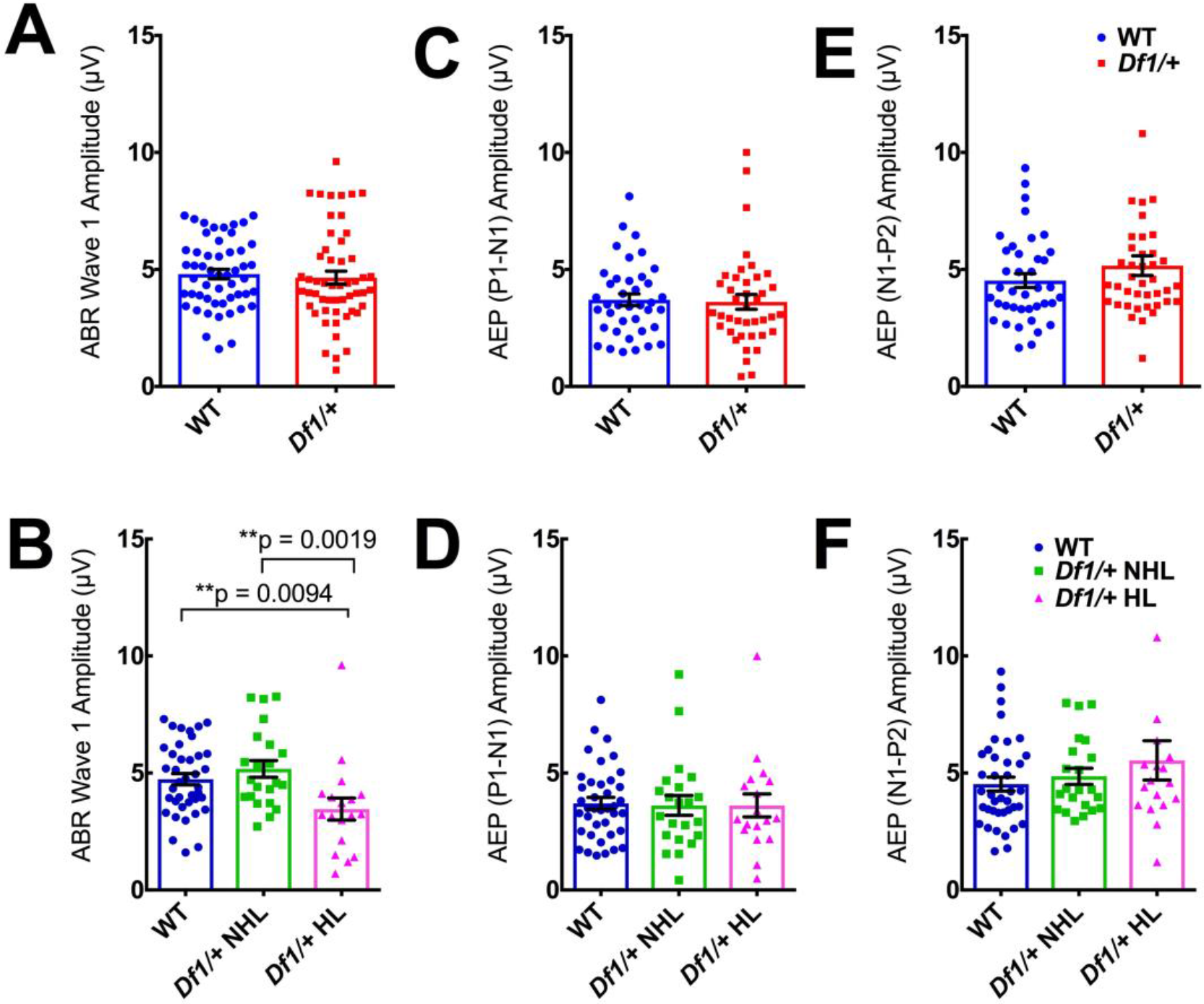
Reductions in ABR wave I amplitude in *Df1*/+ mice with hearing loss are not maintained in cortical AEPs. (A) ABR wave I amplitude to an 80 dB SPL click does not differ between WT and *Df1*/+ mice overall (unpaired *t-*test, p=0.66). (B) However, wave I amplitude is reduced in *Df1*/+ mice with hearing loss relative to either WT mice or *Df1*/+ mice without hearing loss (ordinary one-way ANOVA, F(2,76)=5.55, group difference p=0.0056; Fisher’s LSD, *Df1*/+ HL vs. WT p=0.0094, *Df1*/+ HL vs. *Df1*/+ NHL p=0.0019), while there is no significant difference in ABR wave I amplitude between *Df1*/+ mice without hearing loss and WT mice (*Df1*/+ NHL vs. WT p=0.33). (C-D) AEP P1-N1 amplitude does not differ between WT and *Df1*/+ mice, either overall (C; unpaired *t*-test, p=0.82) or when *Df1*/+ mice with and without hearing loss are considered separately (D; ordinary one-way ANOVA, F(2,76)=0.025, group difference p=0.98). (E-F) No significant differences in AEP N1-P2 amplitude between WT and *Df1*/+ mice, either overall (E; unpaired *t*-test, p=0.22), or when *Df1*/+ mice with and without hearing loss are considered separately (F; ordinary one-way ANOVA, F(2,76)=1.20, group difference p=0.31). Number of mice, number of ABR/AEP recordings, and color conventions as in Figure 2. Bars and error bars indicate mean ± SEM across recordings.

**Figure 4.**
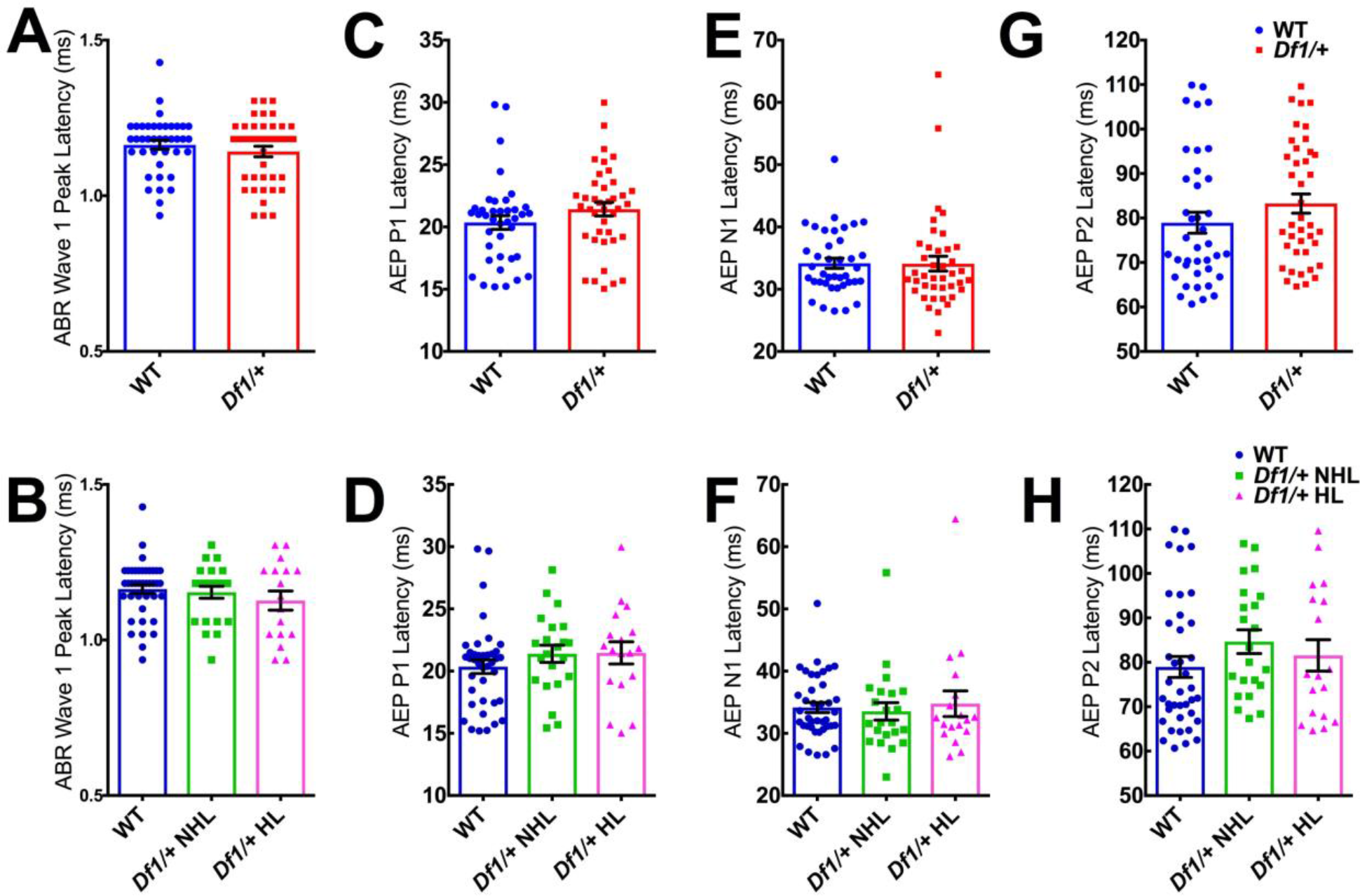
No significant differences between *Df1*/+ and WT mice in ABR wave I latency or AEP P1, N1 or P2 latencies. (A-B) There were no significant differences in ABR wave I peak latency between WT and *Df1*/+ mice overall (A; unpaired *t-*test, p=0.34), nor between WT mice, *Df1*/+ mice without hearing loss, and *Df1*/+ mice with hearing loss (B; one-way ANOVA, F(2,76)=0.81, group difference p=0.45). (C-D) Similarly, there were no significant differences in AEP P1 latency between WT and *Df1*/+ mice overall (C; unpaired *t-*test, p=0.17), nor between WT mice, *Df1*/+ mice without hearing loss, and *Df1*/+ mice with hearing loss (D; one-way ANOVA, F(2,76)=0.94, group difference p=0.39). (E-F) No significant differences in AEP N1 latency between WT and *Df1*/+ mice overall (E; unpaired *t-* test, p=0.97), nor between WT mice, *Df1*/+ mice without hearing loss, and *Df1*/+ mice with hearing loss (F; one-way ANOVA, F(2,76)=0.18, group difference p=0.84). (G-H) No significant differences in AEP P2 latency between WT and *Df1*/+ mice overall (G; unpaired *t-*test, p=0.18), nor between WT mice, *Df1*/+ mice without hearing loss, and *Df1*/+ mice with hearing loss (F; one-way ANOVA, F(2,76)=1.14, group difference p=0.33). Number of mice and number of ABR/AEP recordings as in Figures 2 and 3; plot conventions as in Figure 3.

Hearing loss, defined here as an elevation of the ABR threshold, would be expected to reduce ABR wave I amplitude for a suprathreshold click. As expected, the amplitude of ABR wave I to an 80 dB SPL click was significantly lower in *Df1*/+ mice with hearing loss (HL) than in either *Df1*/+ mice with no hearing loss (NHL) or WT mice (Figure 2C and Figure 3B; ordinary one-way ANOVA, F(2,76)=5.55, group difference p=0.0056; Fisher’s LSD, *Df1*/+ HL vs. NHL p=0.0019; *Df1*/+ HL vs. WT p=0.0094), while there was no significant difference between *Df1*/+ mice without hearing loss and WT animals (*Df1*/+ NHL vs. WT p=0.33), nor between *Df1*/+ and WT mice overall (Figure 2B and Figure 3A; unpaired t-test, p=0.66). More surprisingly, there were no significant differences in AEP wave amplitudes between *Df1*/+ and WT mice, even when *Df1*/+ mice with and without hearing loss were considered separately (Figure 2E,F and Figure 3C-F; WT vs. *Df1*/+ overall, unpaired t-tests, P1-N1: p=0.82, N1-P2: p=0.22; WT vs. *Df1*/+ NHL vs. *Df1*/+ HL, ordinary one-way ANOVAs, P1-N1: F(2,76)=0.025, group difference p=0.98, N1-P2: F(2,76)=1.20, group difference p=0.31). There were also no significant differences in latencies of ABR wave I or AEP waves P1, N1 or P2 between WT and *Df1*/+ animals, either overall or when hearing loss in *Df1*/+ mice was taken into account (Figure 4; unpaired t-tests and ordinary one-way ANOVAs, all comparisons p>0.1). Thus, while ABR wave I amplitude was reduced as expected in *Df1*/+ mice with hearing loss, there were no significant differences in the AEP waves between any of the subgroups, suggesting possible central auditory compensation for reduction in auditory nerve input in *Df1*/+ mice with hearing loss.

### *Df1*/+ mice with hearing loss show elevation of central auditory gain

Given that peripheral hearing loss, including disruption of middle ear function, is known to drive compensatory changes throughout the auditory brain that reduce inhibition and increase “central auditory gain” (Clarkson et al., 2016; Sanes and Kotak, 2011; Takesian et al., 2009; Teichert et al., 2017), we wondered if central auditory gain might differ between *Df1*/+ and WT mice, or between *Df1*/+ mice with and without hearing loss. To determine this, we compared ABR wave I amplitude in each recorded ear to AEP P1-N1 or N1-P2 amplitude recorded over the contralateral cortical hemisphere (number of ear-hemisphere comparisons: 39 WT, 40 *Df1*/+ of which 22 were from *Df1*/+ mice without hearing loss, 18 from *Df1*/+ mice with hearing loss), and we used the ratios of AEP P1-N1 and N1-P2 amplitude to ABR wave I amplitude as measures of central auditory gain in each ABR/AEP recording.

For both the P1-N1 and N1-P2 complexes, central auditory gain was elevated in *Df1*/+ mice with hearing loss (Figure 5). There were no significant differences observed in the ratio of either AEP P1-N1 amplitude or N1-P2 amplitude to ABR wave I amplitude when comparing *Df1*/+ mice overall with WT mice (Figure 5A,C: unpaired t-tests, P1-N1 p=0.36, N1-P2 p=0.084). However, the P1-N1 / wave I ratio was significantly higher in *Df1*/+ mice with hearing loss than in either WT mice or *Df1*/+ mice without hearing loss (Figure 5B; ordinary one-way ANOVA, F(2,76)=4.96, group difference p=0.0094; Fisher’s LSD, *Df1*/+ HL vs. WT p=0.011, *Df1*/+ HL vs. *Df1*/+ NHL p=0.0037), while *Df1*/+ mice without hearing loss were not significantly different from WT animals (WT vs. *Df1*/+ NHL p=0.44). Similarly, the N1-P2 / wave I ratio was significantly higher in *Df1*/+ mice with hearing loss than in either WT mice or *Df1*/+ mice without hearing loss (Figure 5D; ordinary one-way ANOVA, F(2,76)=7.68, group difference p=0.0009; Fisher’s LSD, *Df1*/+ HL vs. WT p=0.0006, *Df1*/+ HL vs. *Df1*/+ NHL p=0.0009), while no significant differences in N1-P2 gain were found between *Df1*/+ mice without hearing loss and WT mice (WT vs. *Df1*/+ NHL p=0.79). These results are consistent with the conclusion that central auditory gain is increased in *Df1*/+ animals with hearing loss.

**Figure 5.**
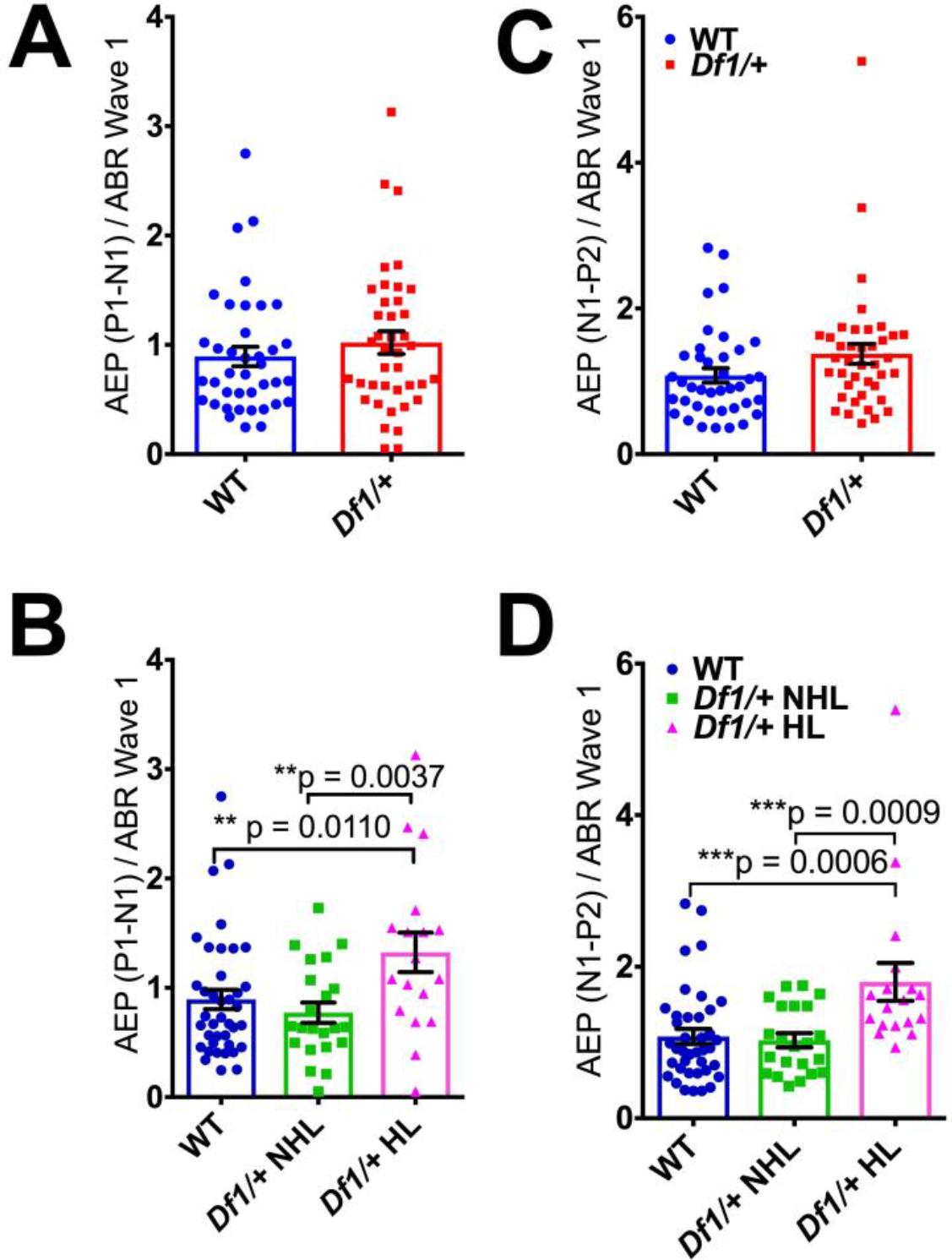
Central auditory gain is elevated in *Df1*/+ mice with hearing loss. (A) The ratio of AEP P1-N1 amplitude to ABR wave I amplitude in ABR/AEP recordings does not differ between WT mice and *Df1*/+ mice overall (unpaired *t-*test, p=0.36). (B) However, this measure of central auditory gain for the P1-N1 complex is significantly elevated in *Df1*/+ mice with hearing loss relative to either WT mice or *Df1*/+ mice without hearing loss (ordinary one-way ANOVA, F(2,76)=4.96, group difference p=0.0094; Fisher’s LSD, *Df1*/+ HL vs. WT p=0.011, *Df1*/+ HL vs. *Df1*/+ NHL p=0.0037, WT vs. *Df1*/+ NHL p=0.44). (C) The ratio of AEP N1-P2 amplitude to ABR wave I amplitude does not differ between WT mice and *Df1*/+ mice overall (unpaired *t-*test, p=0.084). (D) Central auditory gain for the N1-P2 complex is significantly elevated in *Df1*/+ mice with hearing loss relative to either WT mice or *Df1*/+ mice without hearing loss (ordinary one-way ANOVA, F(2,76)=7.68, group difference p=0.0009; Fisher’s LSD, *Df1*/+ HL vs. WT p=0.0006, *Df1*/+ HL vs. *Df1*/+ NHL p=0.0009, WT vs. *Df1*/+ NHL p=0.79). Number of mice and number of ABR/AEP recordings as in Figures 2 and 3; plot conventions as in Figure 3.

### *Df1*/+ mice exhibit abnormal PPI of ASR for prepulse cues with fixed absolute sound level, but not for cues adjusted relative to startle threshold

We next examined potential behavioral influences of the hearing deficit and altered central auditory gain observed in *Df1*/+ mice. The acoustic startle response (ASR) is a reflexive jump evoked by a loud sound, and prepulse inhibition (PPI) of the ASR is the suppression of this jump that occurs when the loud sound is reliably preceded by a quieter but audible cue (the prepulse). Deficits in PPI of the ASR have been reported in the *Df1*/+ mouse and other mouse models of 22q11.2DS (Chun et al., 2014; Didriksen et al., 2017; Paylor and Lindsay, 2006), as well as in humans with 22q11.2DS (Sobin et al., 2004) or schizophrenia (Turetsky et al., 2007). We wondered whether hearing loss and/or central auditory consequences of hearing loss might account for abnormally weak PPI of the ASR in *Df1*/+ mice. To examine this, we measured startle responses and PPI of the ASR in 14 WT mice and 17 *Df1*/+ mice. For testing PPI of the ASR, we chose prepulse sound levels for each mouse that were 5-20 dB below the animal’s behaviorally determined startle response threshold (i.e., below the quietest sound that evoked a significant startle response), thereby adjusting the prepulse cue levels for each animal to be as audible as possible without evoking a startle response. The loud startle-eliciting stimulus, in contrast, was always 95 dB SPL (Willott et al., 1984). All mice included in this analysis exhibited a significant startle response to the 95 dB SPL sound, and no significant startle response to the prepulse cues. Behavioral testing was performed blind to the genotype of the animal (and to its ABR thresholds, which were measured only after behavioral testing was completed).

Startle response functions were determined for each animal by presenting noise bursts of varying sound levels against a 65 dB SPL white noise background. We defined the startle threshold as the lowest sound level that evoked a significant startle response at that sound level and all higher sound levels (Figure 6A,B; see Materials and Methods). We also measured the magnitude of the uncued startle response to the fixed 95 dB SPL noise burst used as the startle-eliciting stimulus in all PPI tests. We found that startle thresholds were significantly higher in *Df1*/+ than in WT mice (Figure 7A; random permutation test, p=0.023), and uncued startle response magnitudes were significantly lower in *Df1*/+ than WT mice (Figure 7C; unpaired t-test, p=0.029). Both of these findings are consistent with the conclusion that hearing loss in *Df1*/+ mice reduces sensitivity to noise bursts used as prepulse cues or startle-eliciting stimuli. We note, however, that the differences between *Df1*/+ and WT mice in startle threshold and uncued startle response magnitude did not reach significance when the *Df1*/+ animals were further subdivided into mice with and without hearing loss (see Figure 7B,D and Figure 7 legend for details).

**Figure 6.**
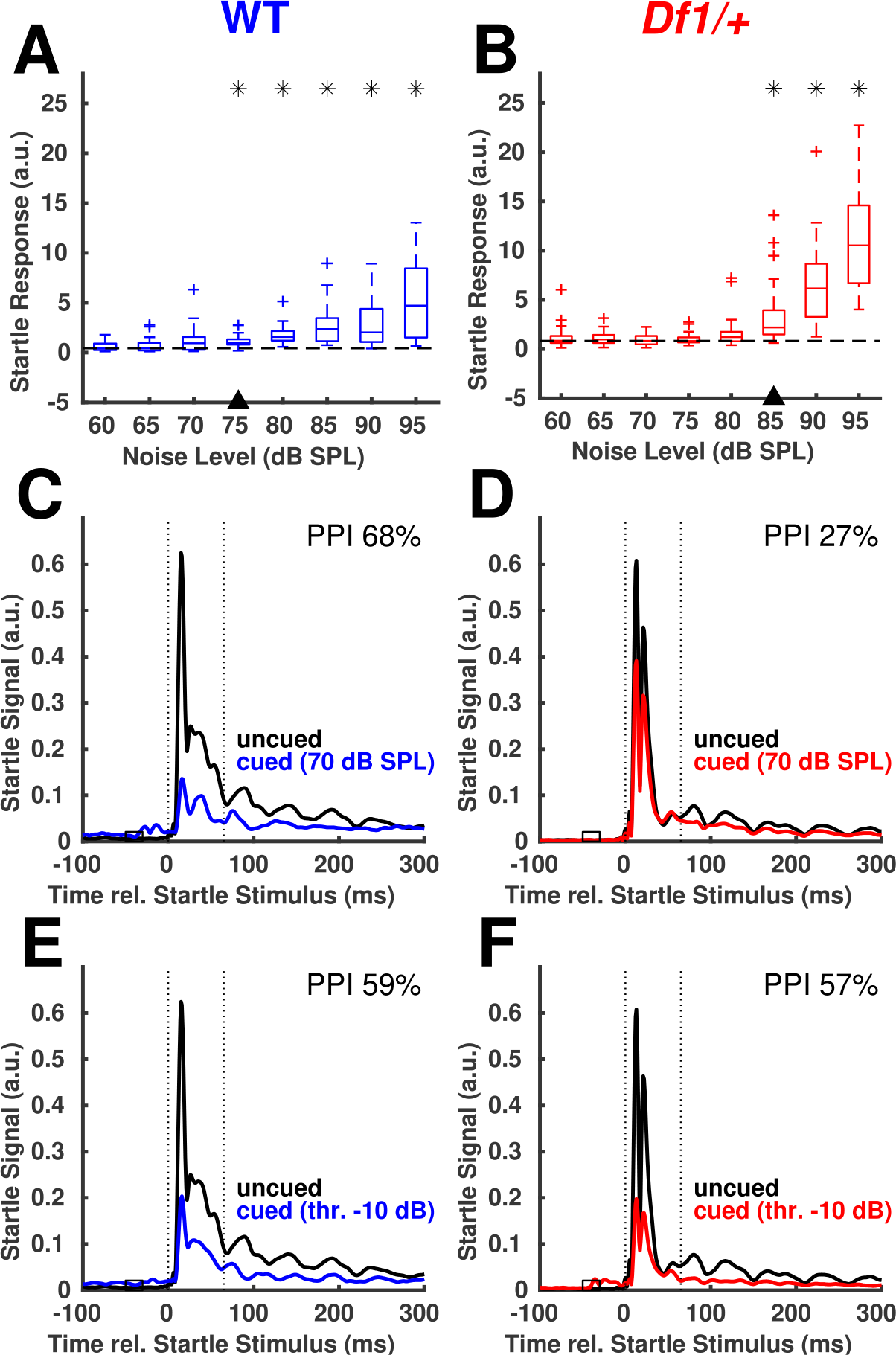
Example acoustic startle responses and prepulse inhibition (PPI) of startle in WT and *Df1*/+ mice. (A) Startle response as a function of noise level for an example WT mouse; boxplots indicate median and quartiles for n=25 trials per condition. Startle response (a.u., arbitrary units) is the sum of startle signal amplitudes (sampled at 1000 samples/sec) over the period 0-65 ms relative to noise burst onset. All noise bursts were presented over a 65 dB SPL continuous white noise background sound. Asterisks indicate noise levels for which the startle response is significantly greater than that evoked by a 60 dB SPL noise burst (Wilcoxon rank-sum test, p<0.01). Triangle on x-axis indicates the startle response threshold, defined as the lowest noise level for which that level and all higher noise levels evoked significant startle responses. Startle response threshold was calculated from data pooled across multiple sessions (as shown) for each mouse. (B) Same as A but for a *Df1*/+ mouse; n=30 trials per condition. Note the elevated startle response (C) Prepulse inhibition of acoustic startle with a 70 dB SPL cue, for the WT mouse. Line plots show startle signal (averaged over n=10 trials per condition) as a function of time relative to startle noise onset. Dotted vertical lines delineate time period used for calculation of startle response. Horizontal open rectangle spanning −50 to −30 ms indicates time of prepulse cue presentation on cued trials. Black trace, uncued trials; colored trace, cued trials. PPI was calculated based on data from a single behavioral session (as shown) and then averaged across repeated sessions for each mouse before population analysis. (D) Same as C but for the *Df1*/+ mouse; n=10 trials per condition. (E) Prepulse inhibition of acoustic startle with cue sound level −10 dB relative to startle threshold, for the WT mouse. (F) Same as E but for the *Df1*/+ mouse. Note that while PPI for a 70 dB SPL prepulse cue is lower for the *Df1*/+ than the WT mouse (C, D), PPI for prepulse cues at −10 dB sound level relative to startle threshold is similar in both mice (E, F).

**Figure 7.**
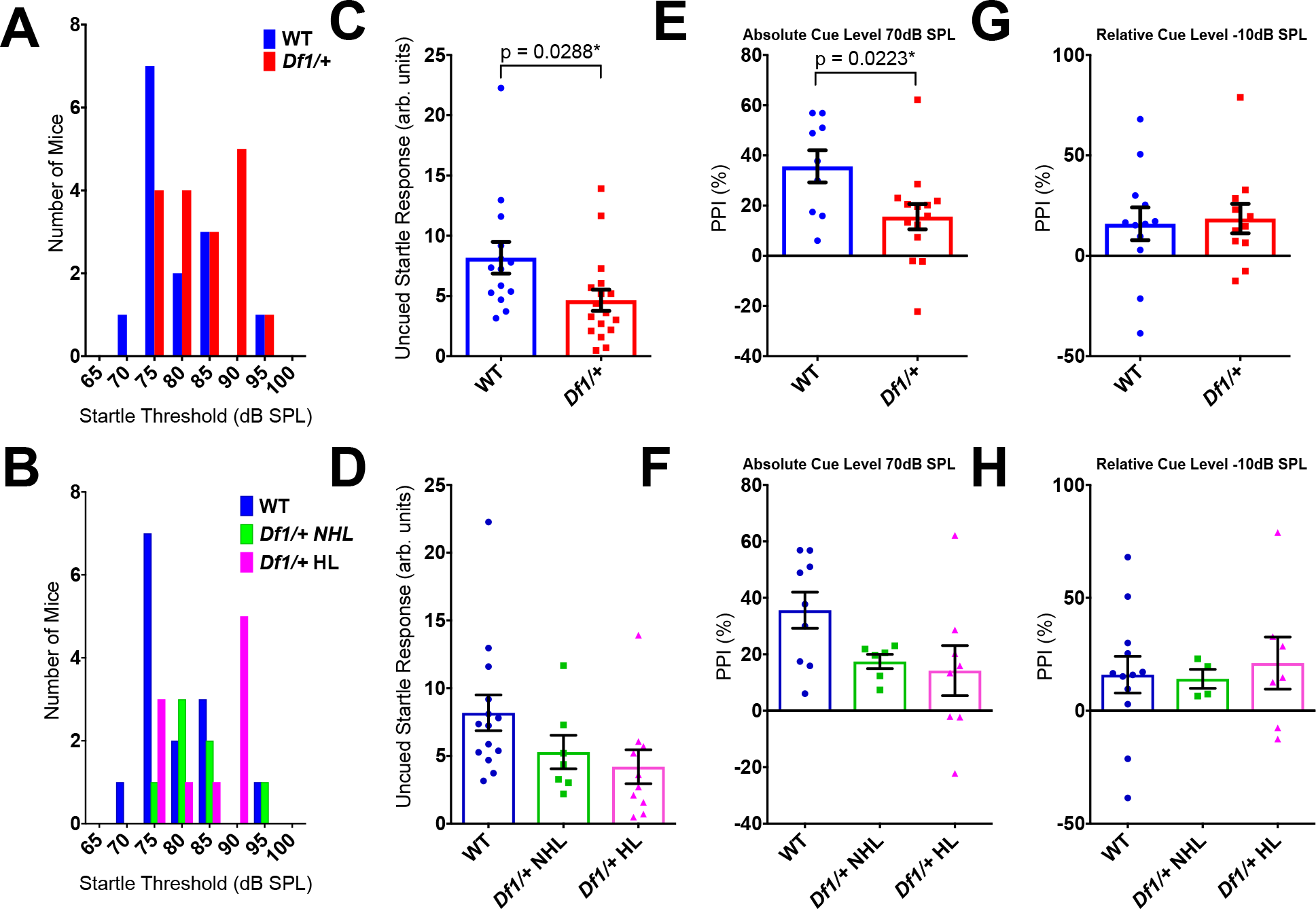
Population analysis of acoustic startle response and prepulse inhibition of startle in WT and *Df1*/+ mice. (A-B) Startle response thresholds are significantly higher in *Df1*/+ mice overall than in WT mice (A; random permutation test, p=0.023), while differences between WT mice, *Df1*/+ mice without hearing loss, and *Df1*/+ mice with hearing loss do not reach significance with correction for multiple testing (B; random permutation tests, p-values, WT vs. *Df1*/+ HL p=0.031, WT vs. *Df1*/+ NHL p=0.078, *Df1*/+ NHL vs. *Df1*/+ HL p=0.30). Number of mice: 14 WT, 17 *Df1*/+ (7 NHL, 10 HL). (C-D) Uncued startle responses to a 95 dB SPL noise burst are significantly lower in *Df1*/+ mice overall than in WT mice (C; unpaired *t-*test, p=0.029), although differences are not significant when groups are further subdivided into WT mice, *Df1*/+ mice without hearing loss, and *Df1*/+ mice with hearing loss (D; ordinary one-way ANOVA, F(2,28)=2.71, group difference p=0.084). Number of mice, same as in A and B. (E-F) Prepulse inhibition with a 70 dB SPL cue, for 9 WT and 14 *Df1*/+ (6 NHL, 8 HL) mice tested at this cue level. PPI is significantly reduced in *Df1*/+ mice overall compared to WT mice (E; unpaired *t-*test, p=0.022). The effect does not reach significance when animal groups are subdivided to compare WT mice and *Df1*/+ mice with and without hearing loss (F; ordinary one-way ANOVA, F(2,20)=2.96, group difference p=0.075). (G-H) PPI for cues with sound level −10 dB relative to startle threshold, for 12 WT and 11 *Df1*/+ (4 NHL, 7 HL) mice tested with this relative cue level. Here, there is no significant difference in PPI either between WT mice and *Df1*/+ mice overall (G; unpaired *t-*test, p=0.82), or between WT mice, *Df1*/+ mice without hearing loss, and *Df1*/+ mice with hearing loss (H; ordinary one-way ANOVA, F(2,20)=0.11, group difference p=0.90).

Several previous studies have reported that PPI of the ASR is reduced in *Df1*/+ mice (Chun et al., 2014; Paylor and Lindsay, 2006). We wondered if PPI of the ASR would still be weaker in *Df1*/+ than WT mice if the sound levels of the prepulse cues were not fixed on an absolute dB SPL scale, but instead adjusted relative to each animal’s startle threshold, to match its sensory salience across animals. The examples of Figure 6 illustrate our approach. After measuring each animal’s startle threshold (Figure 6A,B), we tested PPI of the ASR using at least two different prepulse sound levels, chosen to be between 5 and 20 dB SPL below the animal’s startle threshold. We then compared PPI for prepulse cues with fixed absolute sound level (Figure 6C,D) to PPI when the prepulse cue level was defined relative to startle threshold for each animal (Figure 6E,F). As shown for the example mice in Figure 6, PPI to a fixed 70 dB SPL prepulse cue was 68% in the WT mouse (Figure 6C) but only 27% in the *Df1*/+ mouse (Figure 6D). In contrast, PPI to a prepulse cue with sound level of −10 dB relative to startle threshold was very similar in both mice (Figure 6E,F; 59% in WT mouse, 57% in *Df1*/+ mouse). This finding held across the population. PPI of the ASR was significantly weaker in *Df1*/+ than WT mice tested with a fixed 70 dB SPL prepulse cue (Figure 7E; unpaired t-test, p=0.022; number of mice: 9 WT, 14 *Df1*/+), but there were no significant differences in PPI between *Df1*/+ and WT animals when the cue level was adjusted to be −10 dB relative to startle threshold (Figure 7G; unpaired t-test, p=0.82; number of mice: 11 WT, 12 *Df1*/+). Similar trends but no significant group differences were found when the groups were further subdivided to compare WT mice and *Df1*/+ mice with and without hearing loss (see Figure 7F,H and Figure 7 legend for details). These results suggest that *Df1*/+ mice do not exhibit reduced PPI of acoustic startle compared to WT mice when the prepulse cue level is adjusted relative to startle threshold for each animal.

### *Df1*/+ mice with hearing loss show reduced density of PV+ but not NeuN+ cells in auditory cortex

After examining physiological and behavioral effects of hearing loss in *Df1*/+ mice, we investigated potential neuroanatomical correlates. Abnormalities in parvalbumin-positive (PV+) interneuron networks are a common finding in animal models of schizophrenia (see Lewis, Curley, Glausier, & Volk, 2012 for a review), and alterations in PV+ interneuron activity have previously been linked both to changes in central auditory gain (Resnik and Polley, 2017) and changes in PPI of acoustic startle (Aizenberg et al., 2015). We wondered whether PV+ interneuron density in auditory cortex might be abnormal in *Df1*/+ mice relative to WT mice, and if so, how these abnormalities might relate to hearing loss. To compare PV+ interneuron density to density of neurons overall, we performed immunohistochemical staining for PV and NeuN (a pan-neuronal marker) in coronal brain sections through the auditory cortex (Figure 8A and Figure 4A) in 26 WT mice and 24 *Df1*/+ mice. The final data set included images from auditory cortex in 66 cortical hemispheres (32 WT, 34 *Df1*/+).

**Figure 8.**
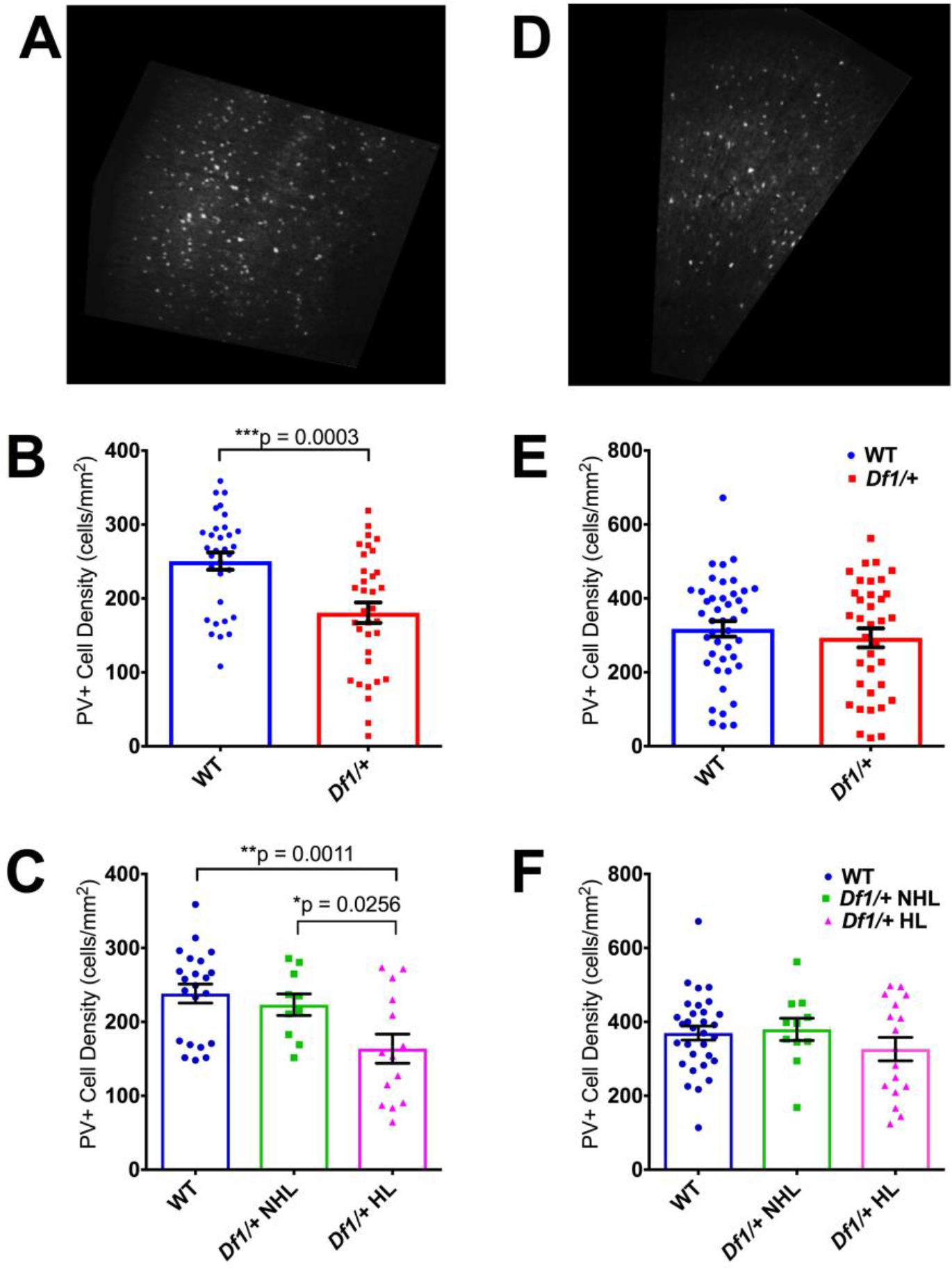
PV+ cell density is reduced in the auditory cortex but not the motor cortex of *Df1*/+ mice. (A) Example confocal image of a coronal section through primary auditory cortex (A1) stained with an antibody against the inhibitory interneuron marker parvalbumin (PV). (B) PV+ cell density in A1 was significantly reduced in *Df1*/+ mice compared to WT mice (unpaired *t-*test, p=0.00030). (C) PV+ cell density was also significantly reduced in *Df1*/+ mice with hearing loss compared to either WT mice or *Df1*/+ mice without hearing loss, but there was no significant difference between WT mice and *Df1*/+ mice without hearing loss (ordinary one-way ANOVA, F(2,43)=6.37, group difference p=0.0038; Fisher’s LSD, *Df1*/+ HL vs. WT p=0.0011, *Df1*/+ HL vs. *Df1*/+ NHL p=0.026, WT vs. *Df1*/+ NHL p=0.53). (D) Example PV-immunostained coronal section through secondary motor cortex (M2). (E) In M2, there was no significant difference in PV+ cell density between *Df1*/+ and WT mice (unpaired *t-*test, p=0.46). (F) PV+ cell density in M2 also did not differ between groups when comparing WT mice, *Df1*/+ mice without hearing loss, and *Df1*/+ mice with hearing loss (ordinary one-way ANOVA, F(2,56)=1.04, group difference p=0.36). See Table 1 for numbers of hemispheres (and mice) in each comparison.

PV+ interneuron density in A1 was significantly lower in *Df1*/+ than WT mice (Figure 8B; unpaired t-test, p=0.00030). Moreover, PV+ cell density was significantly reduced in *Df1*/+ mice with hearing loss compared to either WT mice or *Df1*/+ mice without hearing loss (Figure 8C; ordinary one-way ANOVA, F(2,43)=6.37, group difference p=0.0038; Fisher’s LSD, *Df1*/+ HL vs. WT p=0.0011, *Df1*/+ HL vs. *Df1*/+ NHL p=0.026), while *Df1*/+ mice without hearing loss did not differ from WT animals (*Df1*/+ NHL vs. WT p=0.53). In contrast, there were no significant differences in NeuN+ cell density in A1 between WT and *Df1*/+ animals overall (Figure 9B; unpaired t-test, p=0.29), nor between WT mice, *Df1*/+ mice without hearing loss, and *Df1*/+ mice with hearing loss (Figure 9C; ordinary one-way ANOVA, F(2,22)=0.72, group difference p=0.50). These results indicate that PV+ interneuron density in A1 is reduced in *Df1*/+ mice with hearing loss, suggesting that cortical changes might contribute to central auditory compensation for hearing loss in *Df1*/+ animals.

**Figure 9.**
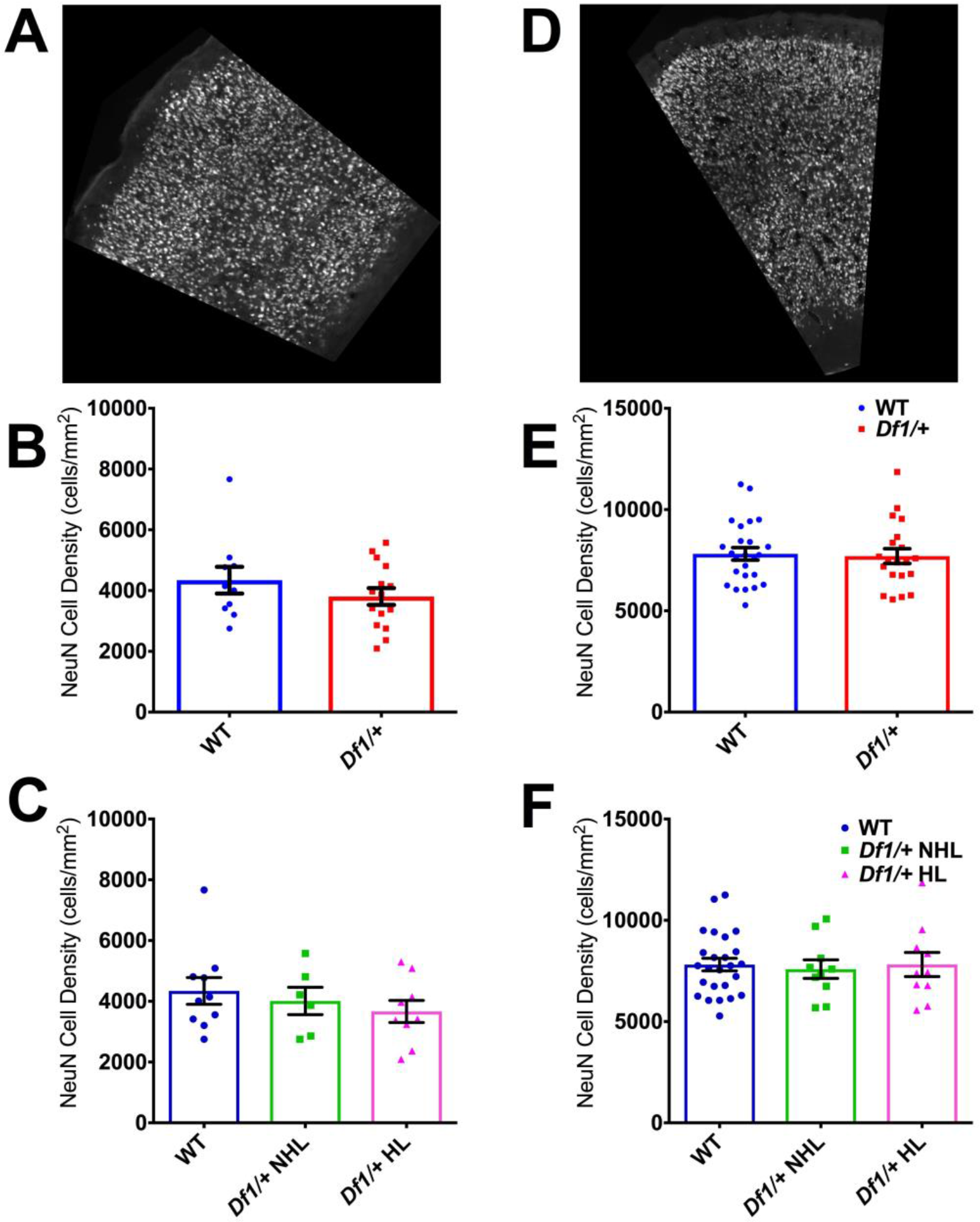
NeuN+ cell density is not reduced in the auditory cortex or the motor cortex of *Df1*/+ mice. (A) Example confocal image of a coronal section through primary auditory cortex (A1) stained with an antibody against the pan-neuronal marker NeuN. (B) NeuN+ cell density in A1 was not significantly different in *Df1*/+ mice compared to WT mice (unpaired *t-*test, p=0.29). (C) No significant difference in A1 NeuN+ cell density between WT mice, *Df1*/+ mice without hearing loss, and *Df1*/+ mice with hearing loss (ordinary one-way ANOVA, F(2,22)=0.72, group difference p=0.50). (D) Example NeuN-immunostained coronal section through secondary motor cortex (M2). (E) In M2, there was no significant difference in NeuN+ cell density between *Df1*/+ and WT mice (unpaired *t-*test, p=0.82). (F) NeuN+ cell density in M2 also did not differ between groups when comparing WT mice, *Df1*/+ mice without hearing loss, and *Df1*/+ mice with hearing loss (ordinary one-way ANOVA, F(2,42)=0.076, group difference p=0.93). See Table 1 for numbers of hemispheres (and mice) in each comparison.

### *Df1*/+ mice do not show abnormalities in PV+ or NeuN+ cell density in M2, nor changes in laminar distributions of PV+ cells in A1

We also wondered if abnormalities in PV+ interneuron density in *Df1*/+ mice might be evident in other cortical areas besides A1. Auditory cortex in mice is reciprocally connected with the secondary motor cortex (M2), and neural activity in M2 is known to modulate auditory cortical processing (Nelson et al., 2013; Schneider et al., 2014). To find out if reductions in PV+ interneuron density observed in A1 of *Df1*/+ mice with hearing loss also occurred in M2, we analyzed PV and NeuN immunostaining in coronal brain sections through frontal cortex (Figure 8D and Figure 9D) in 79 hemispheres (43 WT, 36 *Df1*/+).

PV+ interneuron density in M2 did not differ between WT and *Df1*/+ mice (Figure 8E; unpaired t-test, p=0.46), nor between WT mice, *Df1*/+ mice without hearing loss, and *Df1*/+ mice with hearing loss (Figure 8F; ordinary one-way ANOVA, F(2,56)=1.04, group difference p=0.36). There were also no significant differences between animal groups in NeuN+ cell density in M2, either between WT and *Df1*/+ mice overall or when the Df1/+ animals were further subdivided into those with and without hearing loss (Figure 9E,F; WT vs. *Df1*/+, unpaired t-test, p=0.82; WT vs. *Df1*/+ NHL vs. *Df1*/+ HL, ordinary one-way ANOVA, F(2,42)=0.076, group difference p=0.93). These results suggest that reductions in PV+ interneuron density observed in *Df1*/+ mice with hearing loss may be specific to the auditory cortex.

Previous studies have reported aberrant inhibitory interneuron distributions in the frontal cortex in another mouse model of 22q11.2DS, the LgDel mouse (Meechan et al., 2009). We therefore wondered whether laminar distributions of PV+ interneurons might be altered in A1 or M2 of *Df1*/+ mice. To find out, we quantified the laminar depth of cell centroids along the pia-to-white-matter axis in both areas, normalizing the depth measures by cortical thickness. We then discretized cell counts into 5, 10, or 20 proportional depth bins to analyze effects of genotype or hearing loss on laminar distribution of cells. We found no significant differences in the laminar distribution of PV+ interneurons in *Df1*/+ mice compared to their WT littermates in either A1 or M2 (Figure 10A,C; two-way ANOVA, A1 WT vs. *Df1/*+, F(4,164)=0.25, p=0.91, M2 WT vs. *Df1*/+ mice F(4,212)=1.13, p=0.34). When we compared WT mice and *Df1*/+ mice with and without hearing loss, we again found no significant differences in laminar distribution of PV+ cells in A1, while a weak effect of hearing loss was observed in M2 when using 5 but not 10 or 20 proportional depth bins (Figure 10B,D; two-way ANOVA, A1 WT vs. *Df1*/+ NHL vs. *Df1*/+ HL, F(8,160)=0.22, p=0.99, M2 WT vs. *Df1*/+ NHL vs. *Df1*/+ HL mice F(8,208)=2.24, p=0.026). Thus, we found minimal evidence for abnormalities in laminar distributions of PV+ interneurons in either A1 or M2 of *Df1*/+ mice, regardless of hearing loss.

**Figure 10.**
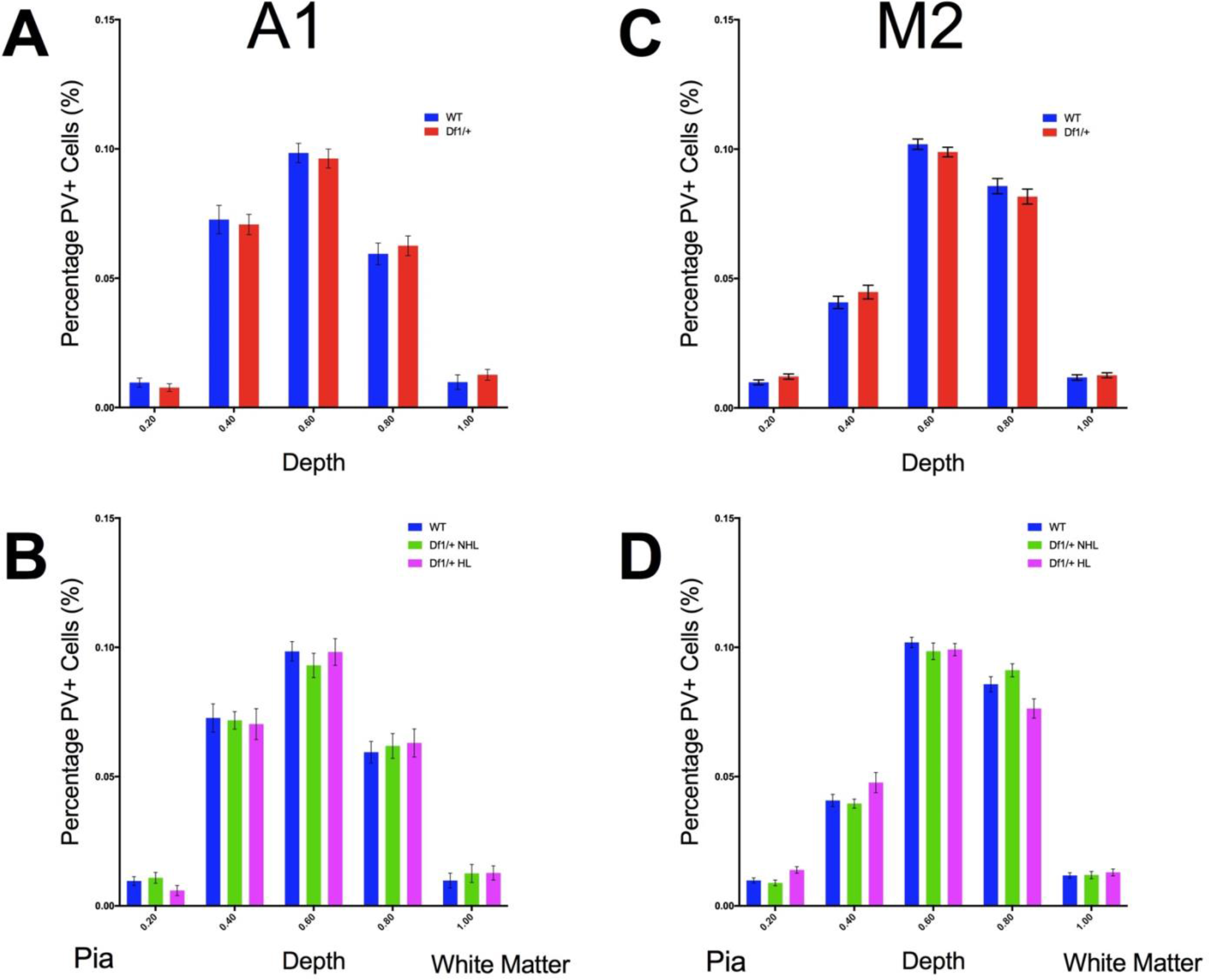
*Df1*/+ mice with hearing loss have minimal differences in laminar distribution of PV+ cells in A1 or M2. Laminar distribution of PV+ cells was analyzed in 5 bins representing equal proportional depths in the cortex from pia to white matter. Error bars indicate SEM across hemispheres; see Table 1 for number of hemispheres and mice. (A,B) PV+ cell laminar distribution in A1, compared between WT and *Df1*/+ mice (A) or between WT mice, *Df1*/+ mice without hearing loss, and *Df1*/+ mice with hearing loss (B). A two-way ANOVA revealed no significant interactions between genotype and bin depth (WT vs. *Df1/+,* F(4,164)=0.25, p=0.91; WT vs. *Df1*/+ NHL vs. *Df1*/+ HL, F(8,160)=0.22, p=0.99). (C,D) PV+ cell laminar distribution in M2, compared between WT and *Df1*/+ mice (C) or between WT mice and *Df1*/+ mice with or without hearing loss (D). Two-way ANOVA revealed no significant interaction between genotype and bin depth for WT vs. *Df1*/+ mice (F(4,212)=1.13, p=0.34), but a weak interaction for WT vs. *Df1*/+ NHL vs. *Df1*/+ HL mice (F(8,208)=2.24, p=0.026). Post hoc Tukey’s multiple comparisons test identified the significant difference as arising from the 0.8 bin, corresponding to depths 0.6-0.8 of the total distance from pia to white matter (WT vs. *Df1*/+ HL p=0.011, *Df1*/+ NHL vs. *Df1*/+ HL p=0. 0022). However, there was no significant interaction between genotype and bin depth nor any single bin with significant difference between groups when the same analysis was performed with 10 or 20 equal-proportion depth bins (all p>0.05).

## DISCUSSION

### Hearing loss in mouse models of 22q11.2DS

Our data confirm that *Df1*/+ mice have high rates of hearing loss, replicating both qualitatively and quantitatively results reported by Fuchs et al. (2013) for a different cohort of animals. Like Fuchs et al. (2013), we found that approximately 60% of *Df1*/+ mice have elevated ABR thresholds in one or both ears; that the incidence of hearing loss is similar in male and female *Df1*/+ mice; and that hearing loss in *Df1*/+ mice can be either monaural or binaural. Fuchs et al. (2013) also showed that hearing loss in *Df1*/+ mice correlates with otitis media (middle ear inflammation). Subsequently, Fuchs et al. (2015) demonstrated that otitis media in *Df1*/+ mice is associated with a bilateral or monolateral defect in the levator veli palatini muscle lining the Eustachian tube, which affects drainage of middle ear effusion. This muscle defect arises early in myogenesis and appears to be caused by haploinsufficiency of the gene *Tbx1* (Fuchs et al., 2015), which has also been linked to other middle and inner ear abnormalities (Liao et al., 2004). Thus, any mouse model of 22q11.2DS with heterozygous deletion of *Tbx1* may be susceptible to otitis media and hearing loss. The *Tbx1* gene lies within the minimum human 22q11.2DS deletion region, and susceptibility to otitis media and hearing loss has been widely reported in human 22q11.2DS patients (see Verheij et al., 2017 for a review).

There are, however, some discrepant results in the literature on hearing loss in mouse models of 22q11.2DS. For example, Paylor et al. (2006) tested frequency-dependent distortion-product otoacoustic emissions (DPOAEs) in 6 *Df1*/+ and 6 WT mice, and found no significant differences in DPOAE thresholds between the groups (see their Supplementary Figure 6). DPOAE thresholds reflect outer hair sensitivity, not auditory nerve activity, but conductive hearing loss (e.g. from otitis media) should increase DPOAE as well as ABR thresholds (Qin et al., 2010). A more direct methodological analogy to Fuchs et al. (2013) and the present study comes from a recent characterisation of a new congenic mouse model of the minimum human 22q11.2DS deletion, the *Df(h22q11)*/+ mouse, which like the *Df1*/+ mouse is heterozygous for deletion of *Tbx1* (Didriksen et al., 2017). Didriksen et al. (2017) found no significant differences in click-evoked ABR thresholds between 13 *Df(h22q11)*/+ and 13 WT animals.

One possible explanation for these discrepancies is differences in methodology. For example, (Paylor et al., 2006) and Didriksen et al. (2017) used in-ear speakers or couplers for sound delivery, rather than free-field speakers as in Fuchs et al. (2013) and the present work. While in-ear and free-field sound delivery techniques should produce similar results, hearing loss arising from middle ear dysfunction might not be detected if physical coupling of a sound delivery device to the ear canal enables transmission of sound signals directly to the inner ear via bone or tissue conduction (Steel et al., 1987).

Age differences and genetic differences in the mice might also help to explain the discrepant results. The *Df(h22q11)1*/+ mice that underwent ABR testing in Didriksen et al. (2017) were indicated to be approximately 5 months old (see their Supplementary Figure A2), while the mice used here and in Paylor et al. (2006) were somewhat younger (here, 1.5-4.5 months old, mean 3 months; 2-4 months old in Paylor et al. 2006). It is possible that the accelerated age-related hearing loss characteristic of the C57BL/6 background strain (Hequembourg and Liberman, 2001) might have obscured hearing loss due to middle ear problems in the Didriksen et al. (2017) study. Alternatively (or additionally), genetic differences between *Df(h22q11)1*/+ and *Df1*/+ mice might have affected results of hearing tests, since the *Df(h22q11)1*/+ mice used by Didriksen et al. (2017) have a slightly larger chromosomal deletion affecting an extra 5 genes (*Dgcr2, Tssk2, Tssk1, Mrpl40,* and *Hira*). However, both *Df1*/+ mice and *Df(h22q11)1*/+ mice exhibit the haploinsufficiency for *Tbx1* that is thought to underlie vulnerability to otitis media and hearing loss (Didriksen et al., 2017; Fuchs et al., 2015), and genetically, the *Df1*/+ mice used by Paylor et al. (2006) should have been essentially identical to the *Df1*/+ mice used here.

Microbiological status of the mice might also be a factor, but is rarely discussed in published reports. Opportunistic pathogens that are common in laboratory mouse facilities can increase risk of otitis media in susceptible animals (Bleich et al., 2008). Mice used in the present study were maintained in standard mouse housing facilities; it is possible that the incidence of otitis media and hearing loss in *Df1*/+ mice might be reduced if animals were housed in a super-clean facility. Importantly, however, hearing loss and otitis media have been estimated to affect a majority of human 22q11.2DS patients (Verheij et al., 2017). Therefore, even if it were possible to reduce the incidence of otitis media in *Df1*/+ mice by restricting their microbiological exposure, the resulting animals would be poorer models of the human syndrome. Similar concerns about the translational value of “squeaky-clean” mouse models have recently been raised by many immunologists (Willyard, 2018).

### Central auditory abnormalities in mouse models of 22q11.2DS

We found that the magnitude of the cortical auditory evoked potential to a loud sound was similar in *Df1*/+ and WT mice, even for *Df1*/+ mice with abnormally elevated ABR thresholds. Measures of central auditory gain (e.g., ratios between AEP P1-N1 or N1-P2 amplitude and ABR wave I amplitude) were significantly higher in *Df1*/+ mice with hearing loss than in WT mice or in *Df1*/+ mice without hearing loss. These results suggest a compensatory increase in central auditory gain in *Df1*/+ mice with hearing loss. Interestingly, although Didriksen et al. (2017) did not observe differences in ABR thresholds between *Df(h22q11*)/+ and WT mice, they did find evidence for increased central auditory gain: AEP amplitude increased more steeply with sound loudness in *Df(h22q11*)/+ than WT animals.

Increased central auditory gain in *Df1*/+ mice with hearing loss could arise at multiple stages of the central auditory pathway. Loss of peripheral auditory input drives homeostatic changes throughout the auditory brainstem, midbrain, thalamus and cortex, which typically manifest as reductions in inhibitory synaptic transmission, increased spontaneous activity, and increased gain of sound-evoked responses (Chambers et al., 2016; Eggermont, 2017; Takesian et al., 2012, 2009). Conductive hearing loss – the form of hearing loss caused by otitis media – has been shown to be sufficient to evoke plastic changes throughout the central auditory system (Clarkson et al., 2016; Sinclair et al., 2017; Teichert et al., 2017; Xu et al., 2007), even when it affects only one ear (Keating et al., 2015; Popescu and Polley, 2010). Moreover, conductive hearing loss that occurs transiently during development can produce a reduction in inhibitory synaptic currents in the auditory cortex that persists in adulthood (Mowery et al., 2015; Sanes and Kotak, 2011; Takesian et al., 2012). Thus, otitis media and hearing loss in mouse models of 22q11.2DS would be expected to produce abnormalities throughout the central auditory system, even if the hearing loss occurred only transiently during development.

Our results are consistent with this literature, and suggest a possible re-interpretation of auditory thalamocortical deficits previously reported in mouse models of 22q11.2DS (Chun et al., 2014). Using in vitro electrophysiology and calcium imaging, Chun et al. (2014) observed a reduction in auditory thalamocortical synaptic transmission in the *Df(16)1*/+ mouse (which has the same deletion as the *Df1*/+ mouse). No such abnormality was detected in visual or somatosensory thalamocortical slices, indicating that the deficit was specific to the auditory system. Chun et al. (2014) linked the disruption of auditory thalamocortical synaptic transmission to aberrant elevation of D2 dopamine receptor (*Drd2*) expression in the auditory thalamus, apparently caused by haploinsufficiency of the microRNA-processing gene *Dgcr8*. They argued that *Drd2*-dependent disruption of thalamic inputs to the auditory cortex could be a key pathogenic mechanism underlying auditory hallucinations in schizophrenia.

A different interpretation is that hearing loss could be the proximal cause of the auditory thalamic and thalamocortical abnormalities reported by Chun et al. (2014). The possibility of hearing loss in *Df(16)1*/+ (or *Dgcr8*/+) animals was not explicitly examined in that study. Furthermore, some of the conclusions were based on analysis of small numbers of in vitro recordings, taken from only a few animals (or an unspecified number of animals). Given that 60% of *Df1*/+ mice have hearing loss in one or both ears (Fuchs et al., 2013 and present work), it is possible that results in Chun et al. (2014) could have been affected by the chance inclusion of differing numbers of animals with or without hearing loss in different analyses. In the absence of animal-by-animal controls for hearing loss, it remains unclear whether elevation of *Drd2* expression in auditory thalamus and disruption of auditory thalamocortical transmission are abnormalities specific to 22q11.2DS models, or more general consequences of hearing loss. Notably, hearing loss has been shown to affect dopaminergic function in the central auditory system (Fyk-Kolodziej et al., 2015; Tong et al., 2005).

### Auditory sensorimotor gating in mouse models of 22q11.2DS

Consistent with findings of previous reports (Didriksen et al., 2017; Paylor et al., 2001; Paylor and Lindsay, 2006; Stark et al., 2008), prepulse inhibition of the acoustic startle response was reduced in *Df1*/+ mice relative to WT mice, for prepulse cues with 70 dB SPL absolute sound level. However, there was no significant difference in PPI between *Df1*/+ and WT mice when the prepulse cue sound level was adjusted to be 10 dB below the startle threshold to match its sensory salience across animals. We also found that uncued startle responses to the startle-eliciting stimulus (a fixed 95 dB SPL noise) were weaker in *Df1*/+ than WT mice.

Acoustic startle thresholds and/or PPI can be affected by abnormalities in either auditory or motor circuitry, or both (Swerdlow et al., 2001). While we cannot rule out the possibility that motor control deficits in *Df1*/+ mice (Sumitomo et al., 2018) might also have contributed to the results, our findings are clearly consistent with known effects of hearing loss. Hearing loss reduces auditory nerve input to the cochlear nucleus, which is the first stage of the acoustic startle circuit, and also alters neural activity in the inferior colliculus, which is involved in PPI (Swerdlow et al., 2001; Takesian et al., 2009).

Central auditory compensation for loss of peripheral input might counteract some effects of hearing loss, but such compensation is likely to be less complete for the subcortical auditory structures providing input to the startle and PPI circuits than for cortical responses (Chambers et al., 2016). Therefore, the fact that cortical AEPs appeared normal in *Df1*/+ mice with hearing loss does not mean that subcortical auditory inputs to the startle and PPI circuits were unimpaired. Similarly, a low threshold for acoustic startle did not necessarily imply normal hearing (cf. Paylor et al., 2001; Scott et al., 2018). While the median startle response threshold was higher in *Df1*/+ than WT animals, the range of thresholds (75-95 dB SPL) largely overlapped in WT mice and *Df1*/+ mice with and without hearing loss.

### Parvalbumin-positive cortical interneurons and hearing loss

In immunohistochemical studies, we found that the density of PV+ inhibitory interneurons was significantly reduced in the auditory cortex of *Df1*/+ relative to WT mice. The reductions in PV+ cell density did not arise from an overall change in neuronal density, and were not observed in the frontal region M2 which is interconnected with the auditory cortex (Nelson et al., 2013; Tomescu et al., 2014). Moreover, the auditory cortex abnormality in PV+ cell density arose primarily in *Df1*/+ mice with hearing loss, again suggesting central auditory compensation for loss of peripheral input.

To our knowledge, this is not only the first report of a link between hearing loss and reduced PV+ interneuron density in an animal model of schizophrenia, but also the first indication that hearing loss due to middle ear inflammation may influence PV+ interneurons in the auditory cortex. Previous studies in mice have demonstrated that PV+ interneurons in the auditory cortex can be affected by age-related (sensorineural) hearing loss (Martin del Campo et al., 2012), or by mutations that disrupt both auditory hair cell function and cortical interneuron migration (Libé-Philippot et al., 2017). It has also been shown that conductive hearing loss during development decreases inhibitory synaptic strength in the auditory cortex (Takesian et al., 2012). Our findings raise the additional possibility that conductive hearing loss, either alone or in combination with genetic risk factors for schizophrenia, may lead to reductions in PV+ interneuron density. Further experiments are required to test this hypothesis in other mouse models with conductive hearing loss, and to identify the reasons for the change in PV+ cell labelling (e.g., disruption of PV+ cell migration, increased PV+ cell death, or decreased PV expression).

### Parvalbumin-positive cortical interneurons in mouse models of 22q11.2DS

Previous studies of mouse models of 22q11.2DS have reported abnormalities in PV+ cortical interneurons in other brain areas and at other developmental stages (Meechan et al., 2015, 2009; Piskorowski et al., 2016). In the Lgdel/+ mouse (with deletion of 5 more genes than in the *Df1*/+ mouse), Meechan et al. (2009) found that corticogenesis and subsequent differentiation of the cerebral cortex was disrupted. Laminar distribution of PV+ interneurons was altered in the medial prefrontal cortex at P21, but there was no apparent reduction in the density of PV+ interneurons (although the density of layer 2/3 pyramidal neurons was reduced). In adult *Df1*/+ and WT mice, we found no significant differences in the laminar distribution of PV+ interneurons in A1 and minimal differences in M2, despite a larger sample size (WT n=30, *Df1*/+ n=25 hemispheres) than that used in the Meechan et al. (2009) study. Therefore, while abnormalities in PV+ interneurons appear to be a reliable finding in mouse models of 22q11.2DS, the form of these abnormalities may differ between mouse models, between brain areas, or between developmental time points. Further work is needed to understand these differences, and also to determine their origins.

Decreased PV+ interneuron density in *Df1*/+ mice might produce abnormalities in cortical dynamics that also occur in schizophrenia. Abnormalities in PV+ interneurons, including reductions in PV+ interneuron density, are a common finding in human schizophrenia (Beasley et al., 2002; Hamburg et al., 2016; Hashimoto et al., 2003; Uhlhaas and Singer, 2010) and are thought to contribute to cognitive dysfunction (Lewis, 2014). Optogenetic reductions in PV+ interneuron activity in animal models significantly attenuate the power of cortical gamma oscillations (Cardin et al., 2009; Sohal et al., 2009). Gamma oscillations are associated with active information processing, and are attenuated in power or synchrony both in animal models of schizophrenia and in human schizophrenia patients (Haenschel et al., 2009; Hamm et al., 2017; Uhlhaas and Singer, 2010).

While changes in PV+ immunostaining in cortex do not necessarily imply changes in interneuron activity or cortical gamma oscillations, our results and those of Meechan et al. (2009, 2015) suggest that mouse models of 22q11.2DS might have chronic abnormalities in PV+ interneuron function. Our results further indicate that hearing loss either drives or contributes to the emergence of auditory cortical abnormalities in mouse models of 22q11.2DS. Further studies are required to disentangle the impact of schizophrenia-associated genetic risk factors from the effects of hearing loss on auditory cortical function.

### Implications for schizophrenia research

Disruption of auditory brain function in *Df1*/+ mice has previously been described as a key schizophrenia-relevant pathology (Chun et al., 2014), but the possible contribution of hearing loss was not considered. Our results point to a significant role for hearing loss in promoting auditory brain and behavioral abnormalities in *Df1*/+ mice. Increased central auditory gain and reduced auditory cortical PV+ interneuron density were most pronounced in *Df1*/+ mice with hearing loss, and there were no significant differences between WT mice and *Df1*/+ mice without hearing loss. Moreover, abnormalities in prepulse inhibition of acoustic startle in *Df1*/+ mice were reduced when prepulse cue level was adjusted relative to each animal’s acoustic startle threshold. Further experiments in other mouse models, including WT mice with induced hearing loss, will be required to determine whether these and other central auditory abnormalities in *Df1*/+ mice arise from hearing loss alone or from an interaction between hearing loss and genetic risk factors for schizophrenia.

There is compelling evidence that hearing loss in adulthood increases the risk of psychosis and hallucinations (see Linszen et al., 2016 for a recent review). Moreover, hearing impairment in childhood elevates the risk of developing schizophrenia in adulthood. The mechanisms underlying the association between hearing loss and psychosis remain unknown, but could include not only common etiology but also bottom-up changes in neuronal networks driven by loss of sensory input, and top-down changes in cognition driven by difficulty with social communication (Linszen et al., 2016). In individuals with genetic or other risk factors for schizophrenia – such as 22q11.2DS patients – these consequences of hearing loss might be a critical “second hit” promoting development of hallucinations and other psychotic symptoms. Thus, the *Df1*/+ mouse and other mouse models of 22q11.2DS may help to reveal how hearing loss interacts with other schizophrenia risk factors to produce brain and behavioral abnormalities underlying psychiatric disease.

## Acknowledgements

This work was supported by National Institute of Mental Health Division of Intramural Research Programs ZIA MH002897 to K.H.W., Q,L. and F.Z., UCL-NIMH GPP program to F.Z., and Action on Hearing Loss G77 to J.F.L. The authors declare no conflict of interest.

## REFERENCES

Aggarwal VS, Liao J, Bondarev A, Schimmang T, Lewandoski M, Locker J, Shanske A, Campione M, Morrow BE. 2006. Dissection of Tbx1 and Fgf interactions in mouse models of 22q11DS suggests functional redundancy. Hum Mol Genet 15:3219–3228. doi:10.1093/hmg/ddl399

Aizenberg M, Mwilambwe-Tshilobo L, Briguglio JJ, Natan RG, Geffen MN. 2015. Bidirectional Regulation of Innate and Learned Behaviors That Rely on Frequency Discrimination by Cortical Inhibitory Neurons. PLOS Biol 13:e1002308. doi:10.1371/journal.pbio.1002308

Beasley CL, Zhang ZJ, Patten I, Reynolds GP. 2002. Selective deficits in prefrontal cortical GABAergic neurons in schizophrenia defined by the presence of calcium-binding proteins. Biol Psychiatry 52:708–715. doi:10.1016/s0006-3223(02)01360-4

Bleich A, Kirsch P, Sahly H, Fahey J, Smoczek A, Hedrich HJ, Sundberg JP. 2008. Klebsiella oxytoca: opportunistic infections in laboratory rodents. Lab Anim 42:369–375. doi:10.1258/la.2007.06026e

Cardin JA, Carlen M, Meletis K, Knoblich U, Zhang F, Deisseroth K, Tsai LH, Moore CI. 2009. Driving fast-spiking cells induces gamma rhythm and controls sensory responses. Nature 459:663–667. doi:10.1038/nature08002

Chambers AR, Resnik J, Yuan YS, Whitton JP, Edge AS, Liberman MC, Polley DB. 2016. Central Gain Restores Auditory Processing following Near-Complete Cochlear Denervation. Neuron 89:867–879. doi:10.1016/j.neuron.2015.12.041

Chun S, Westmoreland JJ, Bayazitov IT, Eddins D, Pani AK, Smeyne RJ, Yu J, Blundon JA, Zakharenko SS. 2014. Specific disruption of thalamic inputs to the auditory cortex in schizophrenia models. Science (80-) 344:1178–1182. doi:10.1126/science.1253895

Clarkson C, Antunes FM, Rubio ME. 2016. Conductive Hearing Loss Has Long-Lasting Structural and Molecular Effects on Presynaptic and Postsynaptic Structures of Auditory Nerve Synapses in the Cochlear Nucleus. J Neurosci 36:10214–10227. doi:10.1523/jneurosci.0226-16.2016

Didriksen M, Fejgin K, Nilsson SRO, Birknow MR, Grayton HM, Larsen PH, Lauridsen JB, Nielsen V, Celada P, Santana N, Kallunki P, Christensen K V., Werge TM, Stensbøl TB, Egebjerg J, Gastambide F, Artigas F, Bastlund JF, Nielsen J, Stensbol TB, Egebjerg J, Gastambide F, Artigas F, Bastlund JF, Nielsen J. 2017. Persistent gating deficit and increased sensitivity to NMDA receptor antagonism after puberty in a new mouse model of the human 22q11.2 microdeletion syndrome: a study in male mice. J Psychiatry Neurosci 42:48–58. doi:10.1503/jpn.150381

Drew LJ, Crabtree GW, Markx S, Stark KL, Chaverneff F, Xu B, Mukai J, Fenelon K, Hsu P-KK, Gogos JA, Karayiorgou M. 2011. The 22q11.2 microdeletion: fifteen years of insights into the genetic and neural complexity of psychiatric disorders. Int J Dev Neurosci 29:259–281. doi:10.1016/j.ijdevneu.2010.09.007

Dyce O, McDonald-McGinn D, Kirschner RE, Zackai E, Young K, Jacobs IN. 2002. Otolaryngologic manifestations of the 22q11.2 deletion syndrome. Arch Otolaryngol Neck Surg 128:1408–1412. doi:10.1001/archotol.128.12.1408

Eggermont JJ. 2017. Acquired hearing loss and brain plasticity. Hear Res 343:176–190. doi:10.1016/j.heares.2016.05.008

Fuchs JC, Linden JF, Baldini A, Tucker AS. 2015. A defect in early myogenesis causes Otitis media in two mouse models of 22q11.2 Deletion Syndrome. Hum Mol Genet 24:1869–82. doi:10.1093/hmg/ddu604

Fuchs JC, Zinnamon FA, Taylor RR, Ivins S, Scambler PJ, Forge A, Tucker AS, Linden JF. 2013. Hearing loss in a mouse model of 22q11.2 Deletion Syndrome. PLoS One 8:e80104. doi:10.1371/journal.pone.0080104

Fyk-Kolodziej BE, Shimano T, Gafoor D, Mirza N, Griffith RD, Gong T-W, Holt AG. 2015. Dopamine in the auditory brainstem and midbrain: co-localization with amino acid neurotransmitters and gene expression following cochlear trauma. Front Neuroanat 9:1–17. doi:10.3389/fnana.2015.00088

Haenschel C, Bittner RA, Waltz J, Haertling F, Wibral M, Singer W, Linden DEJ, Rodriguez E. 2009. Cortical Oscillatory Activity Is Critical for Working Memory as Revealed by Deficits in Early-Onset Schizophrenia. J Neurosci 29:9481–9489. doi:10.1523/jneurosci.1428-09.2009

Hamburg H, Trossbach S V., Bader V, Chwiesko C, Kipar A, Sauvage M, Crum WR, Vernon AC, Bidmon HJ, Korth C. 2016. Simultaneous effects on parvalbumin-positive interneuron and dopaminergic system development in a transgenic rat model for sporadic schizophrenia. Sci Rep 6:1–14. doi:10.1038/srep34946

Hamm JP, Peterka DS, Gogos JA, Yuste R. 2017. Altered Cortical Ensembles in Mouse Models of Schizophrenia. Neuron 94:153–+. doi:10.1016/j.neuron.2017.03.019

Hashimoto T, Volk DW, Eggan SM, Mirnics K, Pierri JN, Sun ZX, Sampson AR, Lewis DA. 2003. Gene expression deficits in a subclass of GABA neurons in the prefrontal cortex of subjects with schizophrenia. J Neurosci 23:6315–6326.

Hequembourg S, Liberman MC. 2001. Spiral ligament pathology: A major aspect of age-related cochlear degeneration in C57BL/6 mice. JARO - J Assoc Res Otolaryngol 2:118–129. doi:10.1007/s101620010075

Keating P, Dahmen JC, King AJ. 2015. Complementary adaptive processes contribute to the developmental plasticity of spatial hearing. Nat Neurosci 18:185–187. doi:10.1038/nn.3914

Lewis DA. 2014. Inhibitory neurons in human cortical circuits: substrate for cognitive dysfunction in schizophrenia. Curr Opin Neurobiol 26:22–26. doi:10.1016/j.conb.2013.11.003

Liao J, Kochilas L, Nowotschin S, Arnold JS, Aggarwal VS, Epstein JA, Brown MC, Adams J, Morrow BE. 2004. Full spectrum of malformations in velo-cardio-facial syndrome/DiGeorge syndrome mouse models by altering Tbx1 dosage. Hum Mol Genet 13:1577–85. doi:10.1093/hmg/ddh176

Libé-Philippot B, Michel V, Boutet de Monvel J, Le Gal S, Dupont T, Avan P, Métin C, Michalski N, Petit C. 2017. Auditory cortex interneuron development requires cadherins operating hair-cell mechanoelectrical transduction. Proc Natl Acad Sci 114:201703408. doi:10.1073/pnas.1703408114

Lindsay EA, Botta A, Jurecic V, Carattini-Rivera S, Cheah YC, Rosenblatt HM, Bradley A, Baldini A. 1999. Congenital heart disease in mice deficient for the DiGeorge syndrome region. Nature 401:379–383. doi:10.1038/43903

Linszen MMJJ, Brouwer RM, Heringa SM, Sommer IE. 2016. Increased risk of psychosis in patients with hearing impairment: Review and meta-analyses. Neurosci Biobehav Rev 62:1–20. doi:10.1016/j.neubiorev.2015.12.012

Maxwell CR, Liang Y, Weightman BD, Kanes SJ, Abel T, Gur RE, Turetsky BI, Bilker WB, Lenox RH, Siegel SJ. 2004. Effects of Chronic Olanzapine and Haloperidol Differ on the Mouse N1 Auditory Evoked Potential. Neuropsychopharmacology 29:739–746. doi:10.1038/sj.npp.1300376

McDonald-McGinn DM, Sullivan KE, Marino B, Philip N, Swillen A, Vorstman JAS, Zackai EH, Emanuel BS, Vermeesch JR, Morrow BE, Scambler PJ, Bassett AS. 2015. 22q11.2 Deletion Syndrome. Nat Rev Dis Prim 1. doi:10.1038/nrdp.2015.71

Meechan DW, Rutz HLH, Fralish MS, Maynard TM, Rothblat LA, LaMantia A-SS. 2015. Cognitive ability is associated with altered medial frontal cortical circuits in the LgDel mouse model of 22q11.2DS. Cereb Cortex 25:1143–51. doi:10.1093/cercor/bht308

Meechan DW, Tucker ES, Maynard TM, LaMantia A-SS. 2009. Diminished dosage of 22q11 genes disrupts neurogenesis and cortical development in a mouse model of 22q11 deletion DiGeorge syndrome. Proc Natl Acad Sci U S A 106:16434–16439. doi:10.1073/pnas.0905696106

Moore AK, Weible AP, Balmer TS, Trussell LO, Wehr M. 2018. Rapid Rebalancing of Excitation and Inhibition by Cortical Circuitry. Neuron 97:1341–+. doi:10.1016/j.neuron.2018.01.045

Mowery TM, Kotak VC, Sanes DH. 2015. Transient Hearing Loss Within a Critical Period Causes Persistent Changes to Cellular Properties in Adult Auditory Cortex. Cereb Cortex 25:2083–2094. doi:10.1093/cercor/bhu013

Nelson A, Schneider DM, Takatoh J, Sakurai K, Wang F, Mooney R. 2013. A circuit for motor cortical modulation of auditory cortical activity. J Neurosci 33:14342–14353. doi:10.1523/JNEUROSCI.2275-13.2013

Parham K. 1997. Distortion product otoacoustic emissions in the C57BL/6J mouse model of age-related hearing loss. Hear Res 112:216–234. doi:10.1016/s0378-5955(97)00124-x

Paxinos G, Franklin KBJ. 2012. The Mouse Brain in Stereotaxic Coordinates, 4th ed. Gulf Professional Publishing.

Paylor R, Glaser B, Mupo A, Ataliotis P, Spencer C, Sobotka A, Sparks C, Choi C-HH, Oghalai J, Curran S, Murphy KC, Monks S, Williams N, O’Donovan MC, Owen MJ, Scambler PJ, Lindsay E. 2006. Tbx1 haploinsufficiency is linked to behavioral disorders in mice and humans-implications for 22q11 deletion syndrome. Proc Natl Acad Sci U S A 103:7729–34. doi:10.1073/pnas.0600206103

Paylor R, Lindsay E. 2006. Mouse Models of 22q11 Deletion Syndrome. Biol Psychiatry 59:1172–1179. doi:10.1016/j.biopsych.2006.01.018

Paylor R, McIlwain KL, McAninch R, Nellis A, Yuva-Paylor LA, Baldini A, Lindsay EA. 2001. Mice deleted for the DiGeorge/velocardiofacial syndrome region show abnormal sensorimotor gating and learning and memory impairments. Hum Mol Genet 10:2645–2650. doi:10.1093/hmg/10.23.2645

Piskorowski RA, Nasrallah K, Diamantopoulou A, Mukai J, Hassan SI, Siegelbaum SA, Gogos JA, Chevaleyre V. 2016. Age-Dependent Specific Changes in Area CA2 of the Hippocampus and Social Memory Deficit in a Mouse Model of the 22q11.2 Deletion Syndrome. Neuron 89:163–176. doi:10.1016/j.neuron.2015.11.036

Popescu M V., Polley DB. 2010. Monaural Deprivation Disrupts Development of Binaural Selectivity in Auditory Midbrain and Cortex. Neuron 65:718–731. doi:10.1016/j.neuron.2010.02.019

Qin ZB, Wood M, Rosowski JJ. 2010. Measurement of conductive hearing loss in mice. Hear Res 263:93–103. doi:10.1016/j.heares.2009.10.002

Resnik J, Polley DB. 2017. Fast-spiking GABA circuit dynamics in the auditory cortex predict recovery of sensory processing following peripheral nerve damage. Elife 6. doi:10.7554/eLife.21452

Reyes MRT, LeBlanc EM, Bassila MK. 1999. Hearing loss and otitis media in velo-cardio-facial syndrome. Int J Pediatr Otorhinolaryngol 47:227–233. doi:10.1016/s0165-5876(98)00180-3

Sanes DH, Kotak VC. 2011. Developmental plasticity of auditory cortical inhibitory synapses. Hear Res 279:140–148. doi:10.1016/j.heares.2011.03.015

Schneider DM, Nelson A, Mooney R. 2014. A synaptic and circuit basis for corollary discharge in the auditory cortex. Nature 513:189–94. doi:10.1038/nature13724

Scott KE, Schormans AL, Pacoli KY, De Oliveira C, Allman BL, Schmid S. 2018. Altered Auditory Processing, Filtering, and Reactivity in the Cntnap2 Knock-Out Rat Model for Neurodevelopmental Disorders. J Neurosci 38:8588–8604. doi:10.1523/JNEUROSCI.0759-18.2018

Sigurdsson T, Stark KL, Karayiorgou M, Gogos JA, Gordon JA. 2010. Impaired hippocampal-prefrontal synchrony in a genetic mouse model of schizophrenia. Nature 464:763–767. doi:10.1038/nature08855

Sinclair JL, Fischl MJ, Alexandrova O, Hess M, Grothe B, Leibold C, Kopp-Scheinpflug C, Heß M, Grothe B, Leibold C, Kopp-Scheinpflug C. 2017. Sound-Evoked Activity Influences Myelination of Brainstem Axons in the Trapezoid Body. J Neurosci 37:8239–8255. doi:10.1523/jneurosci.3728-16.2017

Sobin C, Kiley-Brabeck K, Karayiorgou M. 2004. Associations between prepulse inhibition and executive visual attention in children with the 22q11 deletion syndrome. Mol Psychiatry 10:553–562. doi:10.1038/sj.mp.4001609

Sohal VS, Zhang F, Yizhar O, Deisseroth K. 2009. Parvalbumin neurons and gamma rhythms enhance cortical circuit performance. Nature 459:698–702. doi:10.1038/nature07991

Sommer IE, Roze CM, Linszen MMJ, Somers M, van Zanten G a. 2014. Hearing loss; the neglected risk factor for psychosis. Schizophr Res 158:266–267. doi:10.1016/j.schres.2014.07.020

Stark KL, Xu B, Bagchi A, Lai W-SS, Liu H, Hsu R, Wan X, Pavlidis P, Mills AA, Karayiorgou M, Gogos JA. 2008. Altered brain microRNA biogenesis contributes to phenotypic deficits in a 22q11-deletion mouse model. Nat Genet 40:751–760. doi:10.1038/ng.138

Steel KP, Moorjani P, Bock GR. 1987. MIXED CONDUCTIVE AND SENSORINEURAL HEARING-LOSS IN LP/J MICE. Hear Res 28:227–236. doi:10.1016/0378-5955(87)90051-7

Sumitomo A, Horike K, Hirai K, Butcher N, Boot E, Sakurai T, Jr FCN, Bassett AS, Sawa A, Tomoda T. 2018. A mouse model of 22q11.2 deletions: Molecular and behavioral signatures of Parkinson’s disease and schizophrenia 1–10. doi:10.1126/sciadv.aar6637

Swerdlow NR, Geyer MA, Braff DL. 2001. Neural circuit regulation of prepulse inhibition of startle in the rat: current knowledge and future challenges. Psychopharmacology (Berl) 156:194–215. doi:10.1007/s002130100799

Takesian AE, Kotak VC, Sanes DH. 2012. Age-dependent effect of hearing loss on cortical inhibitory synapse function. J Neurophysiol 107:937–47. doi:10.1152/jn.00515.2011

Takesian AE, Kotak VC, Sanes DH. 2009. Developmental hearing loss disrupts synaptic inhibition: implications for auditory processing. Futur Neurol 4:331–349. doi:10.2217/FNL.09.5

Teichert M, Liebmann L, Hubner CA, Bolz J. 2017. Homeostatic plasticity and synaptic scaling in the adult mouse auditory cortex. Sci Rep 7:14. doi:10.1038/s41598-017-17711-5

Tomescu MI, Rihs TA, Becker R, Britz J, Custo A, Grouiller F, Schneider M, Debbane M, Eliez S, Michel CM, Debbané M, Eliez S, Michel CM. 2014. Deviant dynamics of EEG resting state pattern in 22q11.2 deletion syndrome adolescents: A vulnerability marker of schizophrenia? Schizophr Res 157:175–181. doi:10.1016/j.schres.2014.05.036

Tong L, Altschuler RA, Holt AG. 2005. Tyrosine hydroxylase in rat auditory midbrain: Distribution and changes following deafness. Hear Res 206:28–41. doi:10.1016/j.heares.2005.03.006

Turetsky BI, Calkins ME, Light GA, Olincy A, Radant AD, Swerdlow NR. 2007. Neurophysiological endophenotypes of schizophrenia: The viability of selected candidate measures. Schizophr Bull 33:69–94. doi:10.1093/schbul/sbl060

Uhlhaas PJ, Singer W. 2010. Abnormal neural oscillations and synchrony in schizophrenia. Nat Rev Neurosci 11:100–113. doi:10.1038/nrn2774

Verheij E, Derks LSM, Stegeman I, Thomeer H. 2017. Prevalence of hearing loss and clinical otologic manifestations in patients with 22q11.2 deletion syndrome: A literature review. Clin Otolaryngol 42:1319–1328. doi:10.1111/coa.12874

Willott JF. 2006. Measurement of the Auditory Brainstem Response (ABR) to Study Auditory Sensitivity in Mice. Curr Protoc Neurosci Chapter 8,:1–12. doi:10.1002/0471142301.ns0821bs34

Willott JF, Kulig J, Satterfield T. 1984. The acoustic startle response in DBA/2 and C57BL/6 mice: relationship to auditory neuronal response properties and hearing impairment. Hear Res 16:161–167. doi:10.1016/0378-5955(84)90005-4

Willyard C. 2018. Send in the germs. Nature 556:16–18.

Xu H, Kotak VC, Sanes DH. 2007. Conductive Hearing Loss Disrupts Synaptic and Spike Adaptation in Developing Auditory Cortex. J Neurosci 27:9417–9426. doi:10.1523/jneurosci.1992-07.2007

Yao JD, Sanes DH. 2018. Developmental deprivation-induced perceptual and cortical processing deficits in awake-behaving animals. Elife 7:30. doi:10.7554/eLife.33891

